# CYRI-A regulates macropinocytic cup maturation and mediates integrin uptake, limiting invasive migration

**DOI:** 10.1101/2020.12.04.411645

**Authors:** Anh Hoang Le, Tamas Yelland, Nikki R. Paul, Loic Fort, Savvas Nikolaou, Shehab Ismail, Laura M. Machesky

**Affiliations:** Cancer Research UK Beatson Institute, Garscube Estate, Switchback Road, Bearsden, Glasgow, G61 1BD, UK; Institute of Cancer Sciences, University of Glasgow, Garscube Estate, Switchback Road, Bearsden, Glasgow, G61 1QH, UK; Department of Cell and Developmental Biology, Medical Research Building III, Vanderbilt University, Nashville, TN, 37240-7935, USA; Department of Chemistry, KU Leuven, Celestijnenlaan 200G, 3001 Heverlee, Belgium

## Abstract

The Scar/WAVE complex is the major driver of actin nucleation at the plasma membrane, resulting in lamellipodia and membrane ruffles. While lamellipodia aid migration, membrane ruffles can generate macropinosomes - cup-like structures - important for nutrient uptake and regulation of cell surface receptor levels. How macropinosomes are formed and the role of the actin machinery in their formation and resolution is still not well understood. Mammalian CYRI-B is a recently described negative regulator of the Scar/WAVE complex by RAC1 sequestration, but its other paralogue, CYRI-A has not been characterised. Here we implicate CYRI-A as a key regulator of macropinosome maturation and integrin internalisation from the cell surface. We find that CYRI-A is recruited to nascent macropinosomes in a transient but distinct burst, downstream of PIP3-mediated RAC1 activation and the initial burst of actin assembly driving cup formation, but upstream of internalisation and RAB5 recruitment to the macropinosome. Together, our data place CYRI-A as a local suppressor of actin dynamics, enabling the resolution of the macropinocytic cup. The failure of CYRI-depleted cells to resolve their macropinocytic cups results in reduced integrin a5b1 internalisation, leading to enhanced spreading, invasive behaviour and anchorage-independent 3D growth. We thus describe a new role for CYRI-A as a highly dynamic regulator of RAC1 activity at macropinosomes, modulating homeostasis of integrin surface presentation, with important functional consequences.

## Introduction

The actin cytoskeleton is a multifaceted network, sculpting membranes for motile protrusions and endocytic uptake and trafficking. The small GTPase RAC1 acts as a regulatory switch at the heart of dynamic actin polymerisation. By activating the Scar/WAVE complex, RAC1 indirectly triggers Arp2/3 complex activation and thus branched actin polymerisation at the membrane-cytoplasm interface. RAC1-induced branched actin underpins lamellipodia, endocytic events and macropinocytic cups (Bloomfield and Kay, 2016; Egami et al., 2014; Hinze, 2018; Ferreira and Boucrot, 2018; Mooren et al., 2012). It induces membrane ruffling, a pre-requisite for macropinocytosis and counteracts membrane tension exerted by the cytoplasmic hydrostatic force to drive membrane curvature and invagination (Carlsson, 2018). However, following RAC1 activation, RAC1 inactivation is also essential for the completion of macropinocytosis (Fujii et al., 2013; Yoshida et al., 2009). Constitutively activating RAC1 by photoactivation led to unresolved membrane invagination until photoactivation was turned off (Fujii et al., 2013). The related process phagocytosis, requires several GAPs in the deactivation of RAC1, but only for particles that are larger than 8μm (Schlam et al., 2015). However, whether these GAPs are involved in macropinocytosis remain unknown. Evidence suggests that a balanced spatial and temporal regulation of RAC1 and its downstream cytoskeletal targets at the site of endocytosis is essential.

Integrins are type-I transmembrane proteins important in cell adhesion and migration. Many of the integrin subunits and complexes along with their endocytic trafficking have been implicated in cancer cell invasion and metastasis (Caswell et al., 2009; Cooper and Giancotti, 2019). In breast cancer, increased surface expression of integrin α5 increases the cell contraction force and invasion (Mierke et al., 2011). In ovarian cancer, integrin α5β1 can be transported towards the invasive front by the WASH complex to increase invasion in 3D (Zech et al., 2011).

Among the trafficking routes such as clathrin- or caveolin-1 dependent endocytosis (Shi and Sottile, 2008), bulk internalisation pathways like macropinocytosis and the CLIC-GEEG pathway direct integrin trafficking and can affect the migration and invasion behaviour of cells (Gu et al., 2011; Moreno-Layseca et al., 2020).

Recently, we identified CYRI proteins as negative regulators of the regulation of the Scar/WAVE complex by RAC1 (Fort et al., 2018). Apart from cell migration, actin polymerisation by the Scar/WAVE complex has also been implicated in uptake of cellular pathogens and nutrients by macropinocytosis and phagocytosis (Humphreys et al., 2016; Veltman et al., 2016). Indeed, CYRI-B has been implicated in reducing *Salmonella* infection (Yuki et al., 2019), regulating T-cell activation (Shang et al., 2018) and potentially acting as a tumour suppressor (Chattaragada et al., 2018). Mammalian cells possess two paralogs of the CYRI protein family named CYRI-A and CYRI-B, which are encoded by the *FAM49A* and *FAM49B* gene, respectively. Nothing is currently known about the cellular function of CYRI-A and its relationship with CYRI-B. Through biochemical analyses and live-cell imaging, we explore the membrane kinetics and the subcellular localisation of CYRIs, with a strong focus on the previously uncharacterised CYRI-A and implicate CYRIs as novel regulators of macropinocytosis. We then connect the function of both CYRI-A and B to the trafficking of integrin α5β1 and their effects on cell migration, cell spreading and cancer cell invasion. We propose CYRI proteins as suppressors of the RAC1-actin signalling axis specifically at the macropinocytic cups to ensure efficient completion and maturation of macropinosomes.

## Results

### CYRI-A suppresses lamellipodial spreading and binds active RAC1

While CYRI-B has been characterised as a RAC1-binding protein that restricts lamellipodia, the role of CYRI-A is unknown. Deletion of CYRI-B promotes a broad lamellipodia phenotype with the enrichment of the Scar/WAVE complex at the leading edge (Fort et al., 2018; Yuki et al., 2019). To query whether CYRI-A expression could rescue the cell spreading phenotype caused by depletion of CYRI-B, we expressed either a FLAG-tagged GFP (control) or CYRI-A-FLAG **(Fig. 1 A**) in control or CYRI-B knockout (KO) COS-7 cell ex3 and ex4.1 and measured the change in cell area and the average intensity of the peripheral Arp2/3 complex. As previously described, CYRI-B KO cells showed enhanced lamellipodia spreading (Fort et al., 2018) **(Fig. 1 A-C**), which was rescued by re-expression of CYRI-A-FLAG (**Fig. 1 D**) which suppressed spreading area (**Fig. 1 B**) and Arp2/3 recruitment to the cell edge (**Fig. 1 C**) back to control levels. Expression of untagged CYRI-A using a bicistronic system also rescued the spread area and Arp2/3 recruitment (**Fig. S1 A**). We thus find that CYRI-A can compensate for the loss of CYRI-B in controlling lamellipodia formation.

**Fig. 1.**
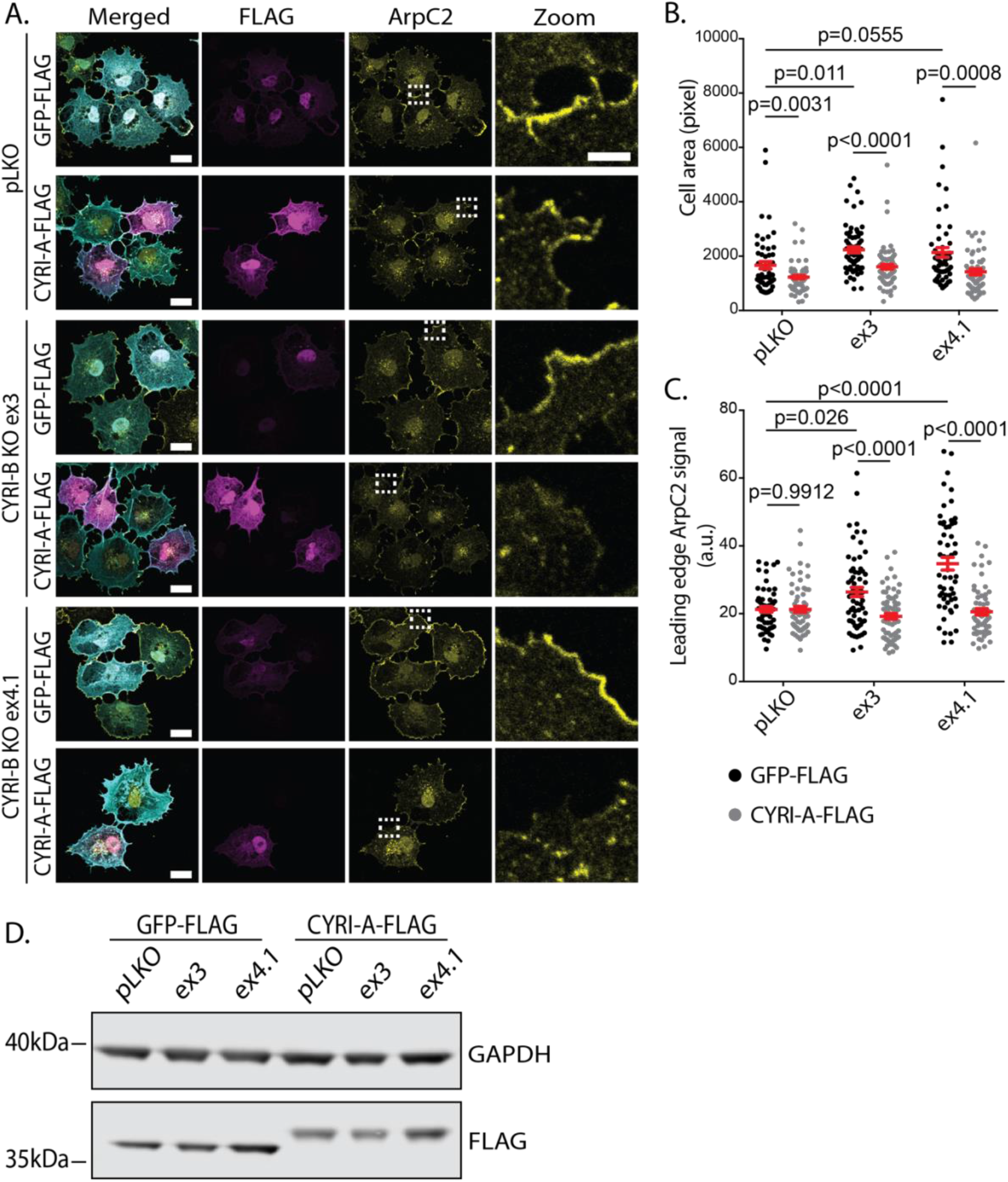
CYRI-A rescues CYRI-B loss phenotype of cell spreading and leading-edge Arp2/3 complex recruitment. A. Immunofluorescence images of COS-7 pLKO (control) and CYRI-B CRISPR KO lines (ex3 or ex4.1) stained for F-actin (phalloidin, cyan), expressing either GFP-FLAG or CYRI-A-FLAG vector (anti-FLAG, magenta) and Arp2/3 complex (anti-ArpC2, yellow), (scale bar = 20μm). Dotted square denotes zooms at right (scale bar = 5μm). Cell spread area and the leading-edge localisation of the Arp2/3 complex reflect the rescue of the CYRI-B deletion phenotype. B-C. Quantification of the cell spread area and Arp2/3 signal at the leading edge in COS-7 cells expressing either GFP-FLAG (black data points) or CYRI-A-FLAG (grey data points). Data from at least 10 cells per experiment and 3 independent experiments. Statistical analyses using two-tailed unpaired t-test. Mean ± SEM. D. Representative Western blot from 1 of the replicates in A shows the relative expression for each construct (GFP-FLAG and CYRI-A-FLAG) in COS-7. GAPDH is used as loading control. Exogenous proteins are detected using anti-FLAG antibody. Molecular size is indicated on the left-hand side.

We next queried whether CYRI-A could interact with active RAC1. Using the I-TASSER protein prediction tool (Roy et al., 2010; Yang et al., 2015; Zhang, 2008) along with the recently solved crystal structures of CYRI-B (Yelland et al., 2020, Kaplan et al., 2020), CYRI-A is predicted to contain an amphipathic N-terminal α-helix connected via a flexible linker to a bundle of 12 α-helices forming the Domain of Unknown Function (or DUF1394 domain) **(Fig. 2 A**). Comparing the amino acid sequence of CYRI-A and CYRI-B shows an 80% sequence identity **(Fig. 2 B**). The two sequences both share a glycine residue at the 2^nd^ position, which is myristoylated in CYRI-B (Green box, **Fig. 2B** and (Fort et al., 2018)) and the two arginine residues at position 159/160 (for CYRI-A) or 160/161 (for CYRI-B) that are essential for active RAC1 binding to CYRI-B (Red box, **Fig. 2B**) and (Fort et al., 2018; Yelland et al., 2020). Using an *in vitro* pulldown assay where GST-tagged RAC1 was immobilised onto beads, we showed that the region of CYRI-A that corresponds to the “RAC1 Binding Domain” or RBD of CYRI-B (Fort et al., 2018) (amino acid 29-319) robustly and consistently interact with active Q61L RAC1 **(Fig. 2 C-D**). To interrogate whether this interaction is direct, surface plasmon resonance (SPR) was performed using purified recombinant maltose binding protein (MBP)-tagged proteins immobilised on an SPR chip. Soluble, untagged constitutively active RAC1 Q61L was titrated in at various concentrations. Surprisingly, CYRI-A interacts with active RAC1 with a Kd of almost one order of magnitude (2.47μM) lower than that of CYRI-B, (18.3μM) similar to the previously reported value (Fort et al., 2018) **(Fig. 2 E-F**). Thus CYRI-A is an active RAC1 interacting protein with a higher affinity for active RAC1 than CYRI-B.

**Fig. 2.**
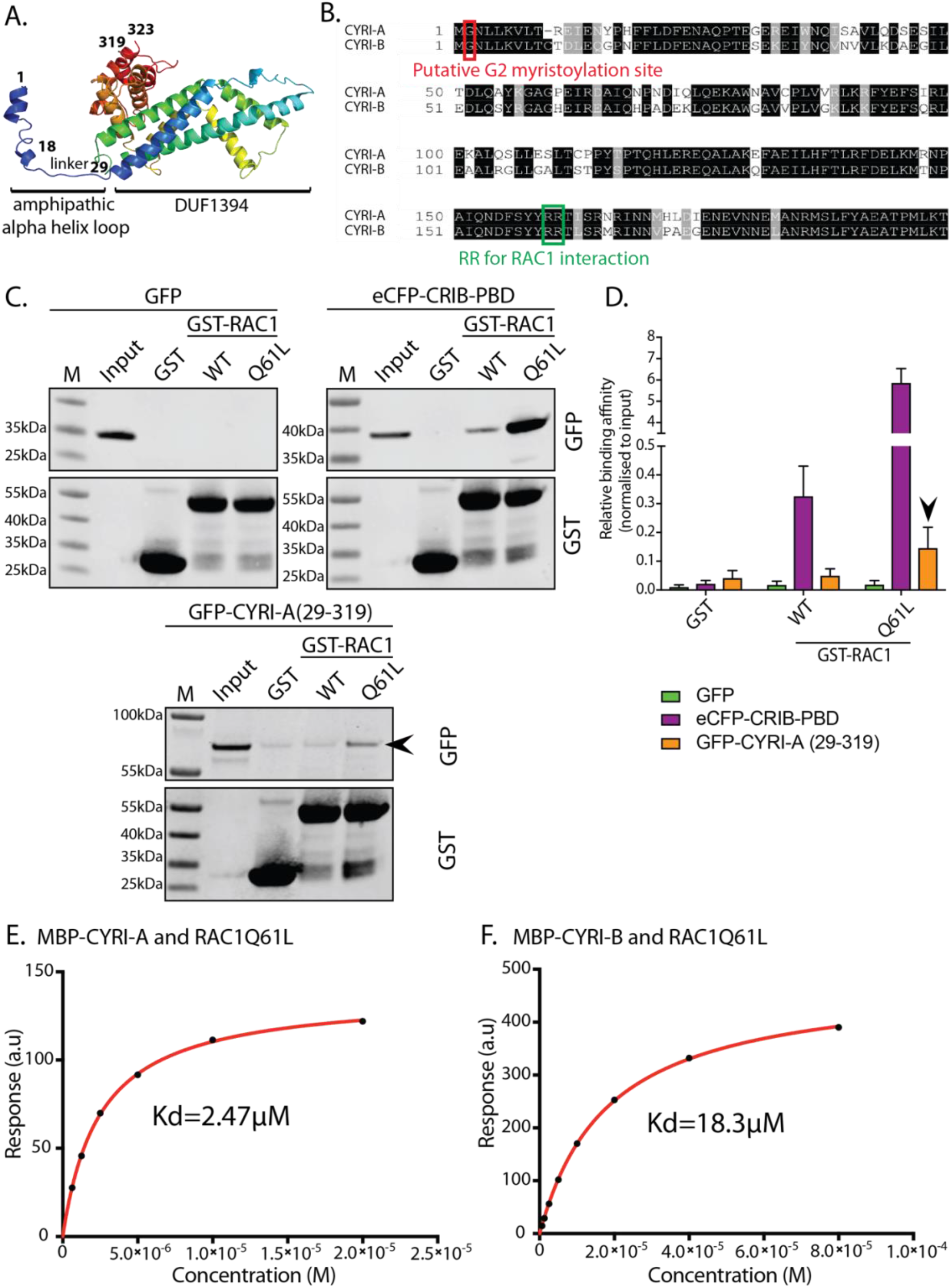
CYRI-A shares large sequence identity with CYRI-B and directly binds active RAC1 with high affinity. A. Predicted 3D fold of CYRI-A by I-TASSER. The protein is composed of one N-terminal amphipathic α-helix connected to a Domain of Unknown Function (DUF) 1394 via a flexible linker. Numbers represent the amino acid position. B. Multiple sequence alignment using Clustal Omega of mouse CYRI-A and CYRI-B shows an 80% sequence identity. Black shades indicate identical residues, grey shades indicate similar residues, white indicates different residues. Highlighted in red is the 2^nd^ Glycine (G2) residues putative myristoylation site. Highlighted in green are two conserved arginine residues (RR) important for mediating active RAC1 binding. C. GST-Trap pulldown assay of GST or GST-RAC1 (WT or Q61L) with lysate from COS-7 cells expressing GFP (negative control), eCFP-CRIB-PBD (positive control) or GFP-CYRI-A (29-319) (RBD). Membranes were blotted with anti-GFP and anti-GST. The black arrowhead denotes the detected interaction between GFP-CYRIA (29-319) and GST-RAC1 Q61L. D. Graph shows the quantification of C) from at least 3 independent experiments. Signals were normalised to the input. Mean ± SEM. Black arrowhead points to RBD and active RAC1 interaction. E-F. Steady state SPR binding curve between MBP-tagged full-length CYRI-A or CYRI-B and increasing concentration of untagged full-length RAC1 Q61L assuming a 1:1 binding model.

### CYRI-A and CYRI-B cooperatively regulate cancer cell spreading and migration

We next sought to characterise the functional relationship of CYRI-A and CYRI-B in a cellular context where both proteins are expressed at comparable levels. HEK293T or COS-7 cells express CYRI-B, but CYRI-A is barely detectable in them (**Fig. S1 B**). Using the EMBL-EBI database, we identified that Ewing’s sarcoma cell lines and particularly A-673 cells as expressing comparable levels of CYRI-A and CYRI-B mRNA. We confirmed the expression of CYRI-A and B protein, and the specificity of the antibody in A-673 cells using siRNA knockdown (KD) or CRISPR-Cas9 knockout (KO) (**Fig. S1 C, Fig. 3 B**). Single knockout of CYRI-A or CYRI-B in A-673 cells resulted in a modest but reproducible effect on the cell shape, with a 10% increase in the number of cells adopting the fast-migrating C-shape, a previously described phenotype (Fort et al., 2018; Yuki et al., 2019) **(Fig. 3 A and C**). However, when both CYRI-A and CYRI-B were simultaneously deleted, this increased to 60% of cells with a C-shape morphology. We observed similar effects in cells treated with siRNA (**Fig. S1 D-E**). DBKO A-673 cells are also accompanied by a 30% increase in the spreading area compared to the control pLKO and single knockout cells. We also measured the migration speed of both CRISPR or siRNA-treated cells on 2D fibronectin substrate and in 3D fibroblast cell-derived matrix (CDM) (Cukierman et al., 2001) **(Fig. 3 D-E, Fig. S1 F-G**). Cells lacking both CYRI isoforms showed a twofold increase in their migration speed (0.5μm/min for DBKO vs 0.2μm/min for pLKO on 2D and 0.2μm/min for DBKO vs 0.1μm/min for pLKO in 3D). There are no obvious differences in motility or morphology between cells lacking just CYRI-A or CYRI-B. Re-introduction of CYRI-A into the DBKO cells reduced the migration speed to basal level **(Fig. 3 F**). Measuring the collective migration behaviour by scratch assay also showed a faster wound closure rate in the DBKO cells compared to either the control pLKO or the single knockout **(Fig. 3 G-H**). DBKO cells proliferate somewhat more slowly than controls (DBKO slope = 0.11, pLKO and single knockout slope = 0.15), making it unlikely that these results are due to changes in proliferation **(Fig. 3 I**). Overall, we find that CYRI-A and CYRI-B have a compensatory role in regulating cell shape and migration in A-673 cells.

**Fig. 3.**
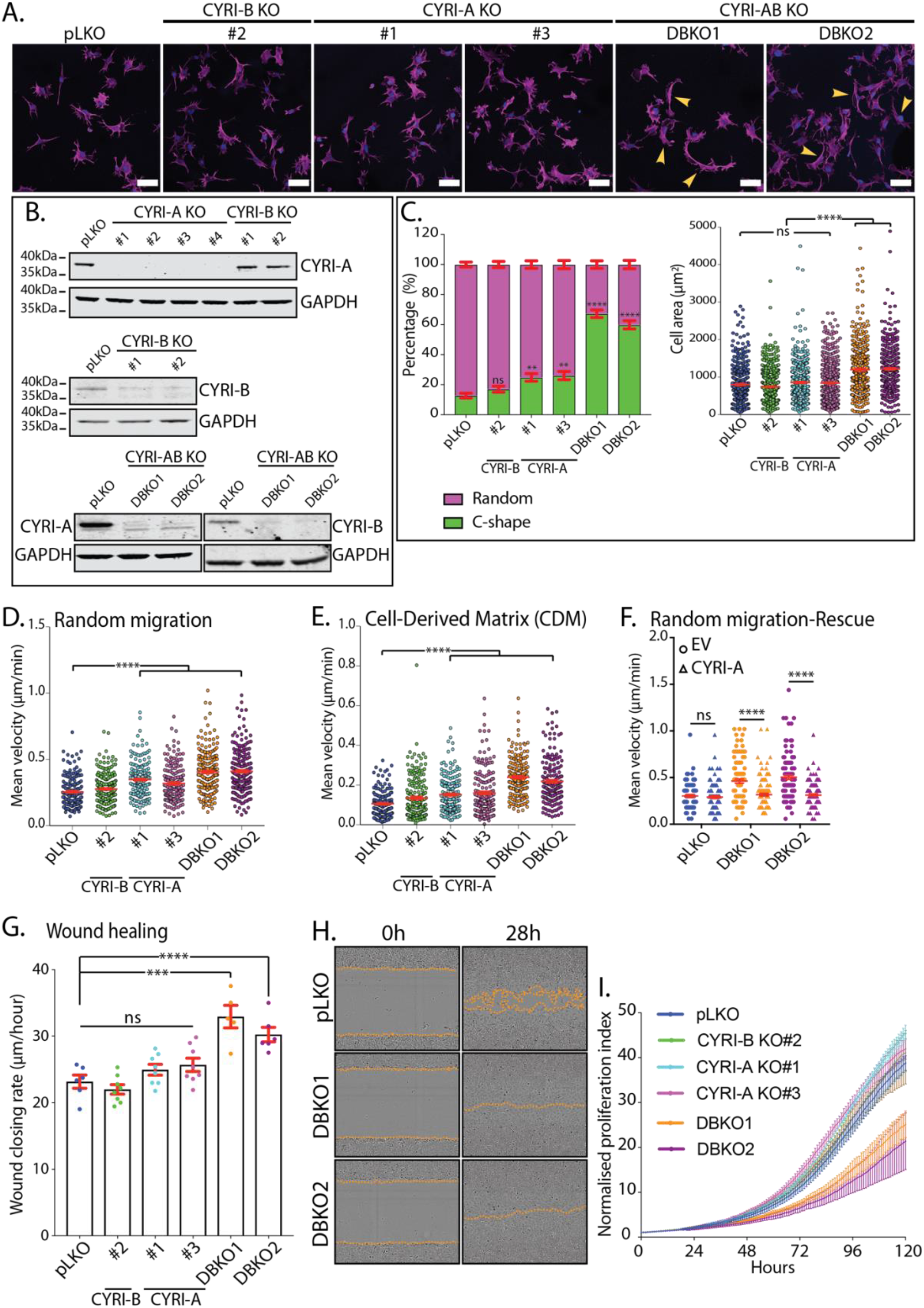
CYRI-A and CYRI-B cooperatively regulate cell shape and migration. All panels show results with A673 cells. A. Immunofluorescence images of control (pLKO), single knockout (CYRI-B = #2,CYRI-A = #1, #3) and double knockout (DBKO1, DBKO2) independent CRISPR cell lines. Cells stained for F-actin (Phalloidin, magenta) and nuclei (DAPI, blue). Yellow arrows indicate C-shaped cells. Scale bar = 50μm. B. Western blots showing the efficiency of single and double knockout of CYRIs. C. Quantification of the cell shape (left) and cell area (right) of A) from at least 50 cells per experiment from 3 independent experiments. D-E. Migration analysis of CYRI CRISPR cells on 2D fibronectin substrate or in 3D cell-derived matrix. F. Effect on cell speed of CYRI-A re-expression in the DBKO cells on 2D fibronectin. G-H. Wound healing assay comparing between the control pLKO, single knockout and double knockout cells. Orange dotted lines highlight the edge of the cell monolayer. I. Proliferation assay using the Incucyte system showing the growth rate of the control pLKO, CYRI-A/B single knockout and double knockouts DBKO1/2. Data from at least 3 independent experiments. Mean ± SEM. ANOVA with Tukey’s multiple comparison test. ns = p>0.05, * p<0.05, ** p<0.01, *** p<0.001, **** p<0.0001.

### CYRI-A is recruited to macropinocytic cups

Our results, combined with previous observations (Fort et al., 2018; Yuki et al., 2019) argue that CYRI proteins interact directly with RAC1 and oppose its activity at the cell leading edge. However, almost nothing is known about the cellular localisation or dynamics of CYRI proteins. Fluorescent tagging of CYRIs has proven difficult, as neither N-nor C-terminal tagging with GFP preserves their dynamic functions (Fort et al, 2018). In an attempt to preserve the N-terminal myristoylation and any role of the N-terminal α-helix, we inserted GFP just after the 16^th^ proline residue of CYRI-A (P16-GFP-CYRI-A) or the 17^th^ proline residue of CYRI-B (P17-GFP-CYRI-B) (**Fig. S1 H**). Both constructs are able to rescue CYRI-B knockout phenotypes in CRISPR COS-7 cells, suggesting that they are functional (**Fig. S1 I-K**). Transiently expressing P16-GFP-CYRI-A results in a striking localisation to many vesicular and occasionally tubular structures **(Fig. 4 A-D, Videos 1-2)**. The size of the vesicles is around 1 μm in diameter, but the length of the tubules can vary from 2 to 8μm. CYRI-A is dynamically localised to vesicles, with resident times of around 50-100s **(Fig. 4 C and G)**.

**Fig. 4.**
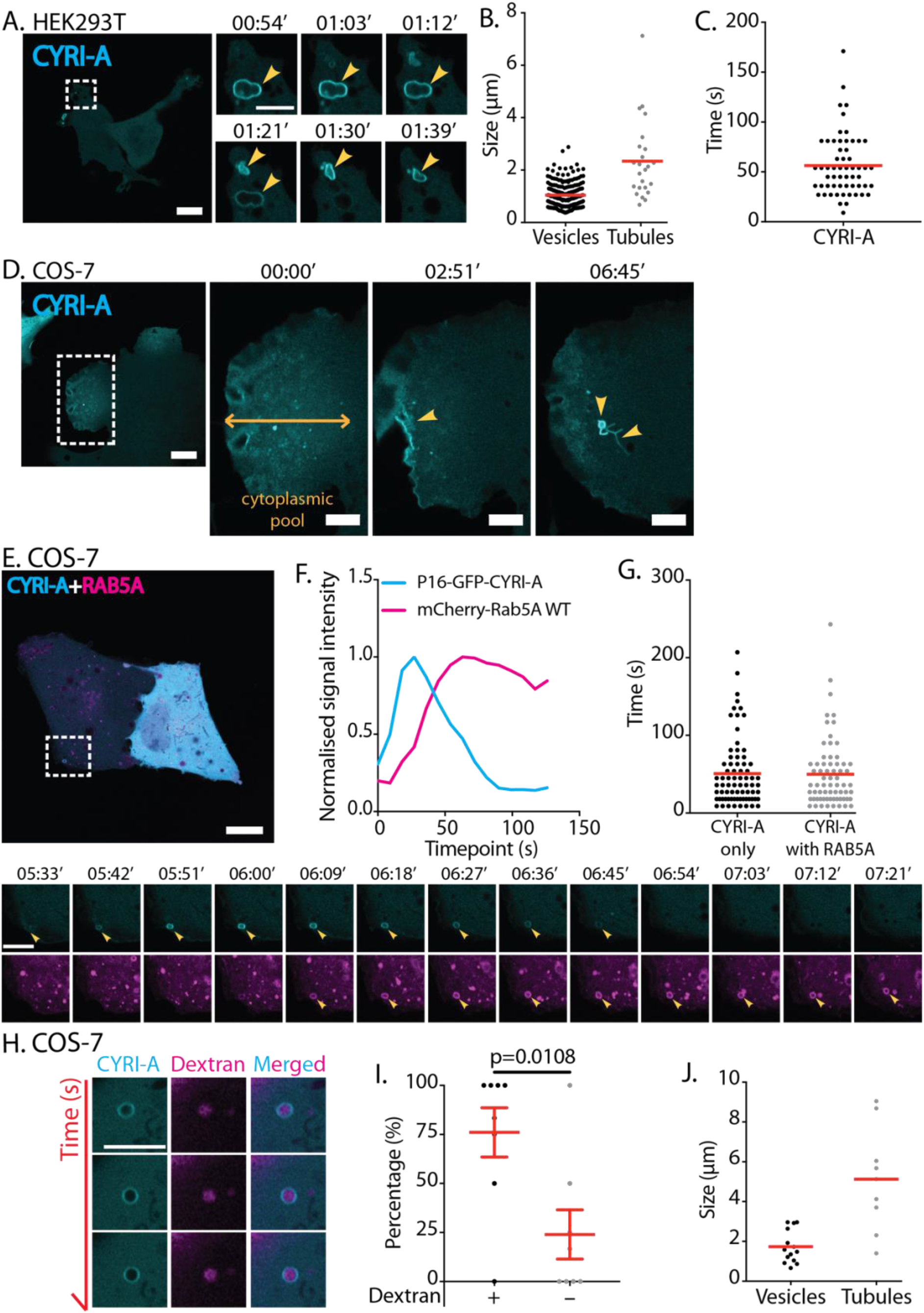
CYRI-A localises to large macropinocytic cup-like structures prior to RAB5A recruitment. A-C. P16-GFP-CYRI-A in HEK293T (scale bar = 20μm) (N=204 events in 18 cells for vesicles and N=24 events in 10 cells for tubules) decorates structures resembling macropinocytic cups (yellow arrows, diameter ranging from 0.4 to 2.9μm, scale bar = 5μm). Tubules with lengths ranging from 0.7 to 7μm. The average lifetime of CYRI-A on cups in HEK293T is about 50s (N=58 events in 5 cells). Red lines represent the average value. See Video 1. D. Still images of COS-7 cells (scale bar = 20μm) expressing P16-GFP-CYRI-A showing the diffused pool of CYRI-A (yellow double arrow) near the leading edge. Dotted square is shown as time sequence on the right (scale bar = 5μm). See Video 2 E. Time-lapse of COS-7 cells expressing P16-GFP-CYRI-A (cyan) and mCherry-RAB5A (magenta). Scale bar = 20μm (full size) or 5μm (zoom). See Video 3. F-G. Quantification showing the time sequence of CYRI-A and RAB5A recruitment to the macropinocytic cups (CYRI-A only N=75 events in 8 cells; CYRI-A with RAB5A N=68 events in 8 cells). H-J. Dextran uptake assay shows CYRI-A decorates dextran-positive vesicles (scale bar = 5μm). Quantification of the percentage of CYRI-A positive vesicles that contain dextran (N=9 cells) and the size of these vesicles and tubules (vesicles N=15 events in 7 cells; tubules N=10 events in 7 cells). Two-tailed unpaired t-test. See Video 4.

Interestingly, we also observe a diffuse localisation of CYRI-A proximal to the cell leading edge in COS-7, which dissipates as CYRI-A is recruited to nascent endocytic cups **(Fig. 4 D**, arrowheads and **Video 2**). In contrast to the dynamic behaviour of CYRI-A, the signal of CYRI-B in COS-7 cells on tubules and vesicles is less dynamic (**Fig. S2 A, Video 21**). The size of CYRI-B positive vesicles is smaller than CYRI-A, with an average of 0.5μm, while tubular structures are more prominent which can reach up to 20μm (**Fig. S2 B**). These observations not only confirm the ability of CYRIs to interact with the cell membrane but also suggest for the first time a potential difference between CYRI-A and CYRI-B in membrane kinetics.

The prominent localisation of CYRIs and the size of these vesicles, in particular CYRI-A, begs the question about the nature of these structures. Interestingly, CYRI-A demonstrates a transient colocalization with mCherry-RAB5A during dual colour live imaging in COS-7 cells **(Fig. 4 E-G, Video 3**). As the vesicles formed, CYRI-A is quickly recruited and remains on the vesicles for around 50s before the RAB5A signal appears. The gradual appearance of RAB5A coincides with the gradual loss of the CYRI-A signal, which takes another 50s **(Fig. 4 G**). RAB5A remains on the vesicles as they are being transported further into the cells. We thus position CYRI-A upstream of RAB5A in internalised vesicles. We also verified this observation in HEK293T cells (**Fig. S2 C-E, Video 22**). On the other hand, CYRI-B remained relatively evenly distributed on vesicles and tubules over these timescales (**Fig. S2 F-G**, **Video 23**). Due to the relatively large size of many of the CYRI-A vesicles, we speculated that they could be macropinosomes (Canton, 2018; Swanson and Watts, 1995). Using fluorescent dextran 70kDa as a marker for fluid-phase uptake, we found that 75% of CYRI-A positive vesicles in COS-7 **(Fig. 4 H-I, Video 4**) and almost 100% in both HEK293T and CHL-1 cells contained dextran (**Fig. S2 H-M, Videos 24-25**). The size of these dextran containing vesicles (0.6-1 μm on average) is within the range of those previously described in **Fig. 4 A-D** and is consistent with macropinosomes (Canton, 2018; Swanson and Watts, 1995). We tested the colocalization of CYRI vesicles with other classical endocytosis pathways including clathrin-mediated endocytosis (CLC15), caveolin-mediated endocytosis (CAV1) and ARF1-dependent endocytosis (ARF1) (**Fig. S3 A-D, Videos 26-29**) but we found no clear colocalization with these markers at any of the observed events. While CYRI-B appears mainly at the plasma membrane, CYRI-A is recruited early in bright flashes to nascent macropinosomes, prior to RAB5A recruitment. Overall, these data highlight a dynamic role for CYRI-A and provide evidence that CYRIs are involved in the early phase of macropinocytosis.

### CYRI-A regulates macropinosome maturation and is in a positive feedback loop with actin dynamics

Actin polymerisation shapes the macropinocytic cup in the early stages (Bloomfield and Kay, 2016; Condon et al., 2018) and CYRI proteins are recruited during RAC1-mediated actin polymerisation in lamellipodia (Fort et al., 2018), so we hypothesized that CYRI-A and actin could coordinate their activities during macropinocytosis. Co-transfecting COS-7 cells with LifeAct-RFP and P16-GFP-CYRI-A shows colocalization between actin and CYRI-A but not with GFP alone **(Fig. 5 A, Videos 5-6**). Tracking the signal intensity between actin and CYRI-A, it was apparent that actin accumulation at the macropinosome preceded CYRI-A accumulation (**Fig. 5 B, Video 7**). Actin is dynamic and drives the early stages of macropinocytic cup formation (Mooren et al., 2012; Bloomfield, 2016; Yoshida, 2009). In COS-7 cells, we often observe the actin signal increases and decreases multiple times around vesicles. Each time the actin signal increases, it is sharply followed by the increase of the CYRI-A signal **(Fig. 5 B, graph**) then both signals decrease, suggesting sequential recruitment. We verified this observation in HEK293T cells **(Fig. 5 C-D, Video 8**). Similarly, CYRI-A signal coincides with the quick decrease in the actin signal and then both signals level off. On average, the actin signal persists on the vesicles for 36s before the CYRI-A signal becomes detectable. CYRI-A and actin coincide for another 54s on average before both decrease, similar to the lifetime of CYRI-A on these vesicles before RAB5A appears (**Fig. S2 E**). To directly compare the effects of CYRIs on the closure and maturation of macropinocytic cups and the resident time of actin on these vesicles, we co-expressed LifeAct-RFP and either GFP alone or the internal GFP-CYRI-A constructs in COS-7 cells that have been depleted of both CYRI-A and CYRI-B using siRNA (DBKD for double knockdown) **(Fig. 5 E-G, Videos 9-10**). As expected, DBKD COS-7 cells are flat with relatively nondynamic lamellipodia. Counting the number of actin-positive large (>0.5μm) vesicles formed shows that GFP-expressing cells form significantly fewer vesicles (~4 vesicles/cell) compared to P16-GFP-CYRI-A rescued cells (~65 vesicles/cell). Measuring the resident time of actin reveals that vesicles recruiting CYRI-A have shorter actin lifetime (~92s) compared to those without CYRI-A (~212s). Indeed, DBKD cells uptake significantly less dextran 70kDa than control cells after 30min of incubation **(Fig. 5 H-I)**. This suggests that CYRI-A suppresses the actin signal around macropinosomes to accelerate their maturation process. To query the possible actin-dependence of CYRI-A localisation, we treated COS-7 and HEK293T cells expressing both CYRI-A and LifeAct with 1 μM of Latrunculin A, which binds to monomeric actin and prevents actin polymerisation (Yarmola et al., 2000) **(Fig. 5 J)**. Within 30s of addition, the actin cytoskeleton collapsed, but CYRI-A vesicles were still frequently observed, even without any obvious membrane ruffles. This suggests that CYRI-A does not directly depend on actin for its localisation. However, Latrunculin A decreases the lifetime of CYRI-A on these large vesicles by approximately 2-fold in both COS-7 and HEK293T cells **(Fig. 5 J)**, suggesting a feedback loop between actin and CYRI-A.

**Fig. 5.**
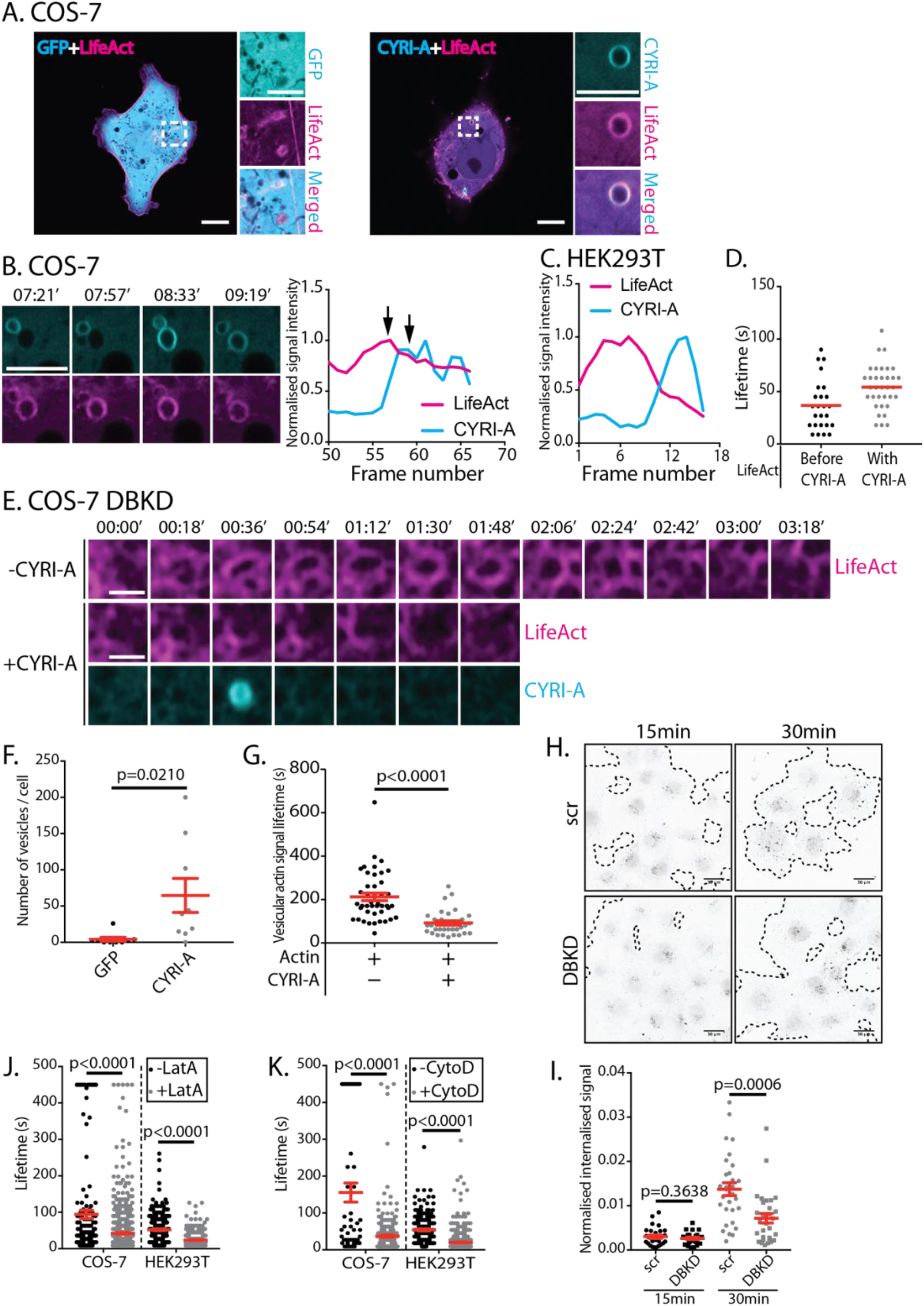
CYRI-A regulates actin accumulation at macropinocytic cups and macropinosome maturation. A. Still images of COS-7 cells expressing either GFP (negative control) or P16-GFP-CYRI-A and LifeAct-RFP. Scale bar = 10μm for full-sized image and 5μm for zooms. See Videos 5-6. B. Time sequence images showing the dynamics of CYRI-A and actin at the macropinosome in COS-7 cells. The graph shows the normalised signal intensities over time. Black arrows show the two peaks of the actin and the CYRI-A signal. Scale bar = 5μm. See also Video 7. C. The line graph shows the normalised signal intensities over time between CYRI-A and LifeAct signal in HEK293T cells. See also Video 8. D. Quantification showing the lifetime of actin before and after CYRI-A is recruited in HEK293T cells (Before CYRI-A: N=25 events in 10 cells, With CYRI-A: N=34 events in 10 cells). Red lines indicate the average value. Scale bar = 10μm for full-sized image and 5μm for zooms. E-G. Time sequence images showing the lifetime of actin on the cups with or without the re-expression of P16-GFP-CYRI-A in CYRI DBKD COS-7 cells. Scale bar = 1μm. Quantification of the number of actin-positive cups between cells with or without P16-GFP-CYRI-A (N=9 cells) (F). Quantification of the lifetime of the actin signal on cups that are negative or positive for CYRI-A signal (Actin alone: N=43 events in 9 cells; Actin with CYRI-A: N=33 events in 8 cells). See also Videos 9-10. H. Macropinocytosis assay in siRNA-treated COS-7 cells. Scr = scramble, DBKD = double knockdown. Black dots are internalised dextran. Black dashed lines indicate the boundary of the cell clusters. Scale bar = 30μm. I. Quantification of the macropinocytic index of H. Data are from at least 10 different fields of view per experiment from a total of 3 independent experiments. Statistical analysis using two-tailed unpaired t-test. Mean ± SEM. J-K. Graphs show the lifetime of CYRI-A on macropinosomes in the absence or presence of Latrunculin A (LatA) or Cytochalasin D (CytoD) in COS-7 and HEK293T cells. Data from 3 independent experiments. Mean ± SEM. Statistical analysis using two-tailed unpaired t-test.

Since Latrunculin A has been shown to activate the Scar/WAVE complex by an unknown mechanism (Millius et al., 2009,Weiner, 2007), which could potentially influence the behaviour of CYRI-A, we reconfirmed our observation using Cytochalasin D, which caps the F-actin and prevents the addition of new actin monomers (Schliwa, 1982). Upon the addition of 1 μM Cytochalasin D, a similar response was observed and the lifetime of CYRI-A on the vesicles was consistently decreased to similar values as in the case of Latrunculin A (**Fig. 5 K**). Overall, these data suggest that CYRI-A is recruited to macropinosomes to oppose positive signals that enhance the actin network such as RAC1 and Scar/WAVE. CYRI-A’s lifetime at the macropinosomes but not its recruitment is actin-dependent. This fits with our hypothesis for CYRI-A as a local inhibitor of actin assembly at macropinosomes, which drives the maturation process.

### CYRI-A regulates actin dynamics through suppressing active RAC1 activity at macropinocytic cups

Direct observation of the spatiotemporal localisation of RAC1 at macropinocytic cups is limited. We thus sought to follow CYRI-A and wild-type RAC1 *in cellulo* to capture their dynamics. Cells overexpressing GFP-tagged RAC1 are broadly spread and display membrane ruffles **(Fig. 6 A, Video 11**). RAC1 signal enriches at the edge of the plasma membrane, especially in ruffles and folds (**Fig. 6 A**). These ruffles fold onto themselves and form macropinosomes (**Fig. 6A**, yellow arrowhead). Importantly, CYRI-A is recruited either slightly after RAC1 or coincident with RAC1 to the cups **(Fig. 6 B-C**). The increase of CYRI-A signal precedes the immediate drop in RAC1 signal, then both signals disappear, as the macropinocytic cup matures. The timing of RAC1 accumulation matches closely to the timing of filamentous actin accumulation on these cups **(Fig. 5 B-D**), suggesting a mechanistic connection between CYRI-A, RAC1 and actin. We next examined the specific localisation of active RAC1 relative to CYRI-A, using the CFP-tagged Pak Binding Domain (PBD) **(Fig. 6 D-E, Video 12**). Active RAC1 (PBD) accumulates as the cups mature and prior to the point where PBD is accompanied by a sharp burst of CYRI-A signal before both dissipate. It is important to emphasize that all of these events happen locally on the macropinocytic cups as they are forming with a high degree of spatiotemporal control. We queried whether CYRI-A’s ability to bind active RAC1 is important for its recruitment to nascent macropinosomes by using a RAC1-binding defective mutant (Yelland et al., 2020) with arginine 159 and 160 mutated to aspartic acid (P16-mCh-CYRI-A-RRDD). We co-transfected COS-7, HEK293T or CHL-1 cells with the wild-type P16-GFP-CYRI-A and either the wild-type or the mutant RRDD construct of P16-mCherry-CYRI-A **(Fig. 6 F-K**, **Videos 13-14**, **Fig. S3 E-F, Videos 30-31**). In all cell types tested, GFP-tagged and mCherry-tagged wild-type CYRI-A show close colocalization at every endocytic event captured. In contrast, mutant CYRI-A did not colocalize with wild-type CYRI-A on any of these endocytic events. Overall, these data strongly suggest that CYRI-A is locally recruited to the macropinocytic cups to suppress actin signalling by responding to the presence of active RAC1.

**Fig. 6.**
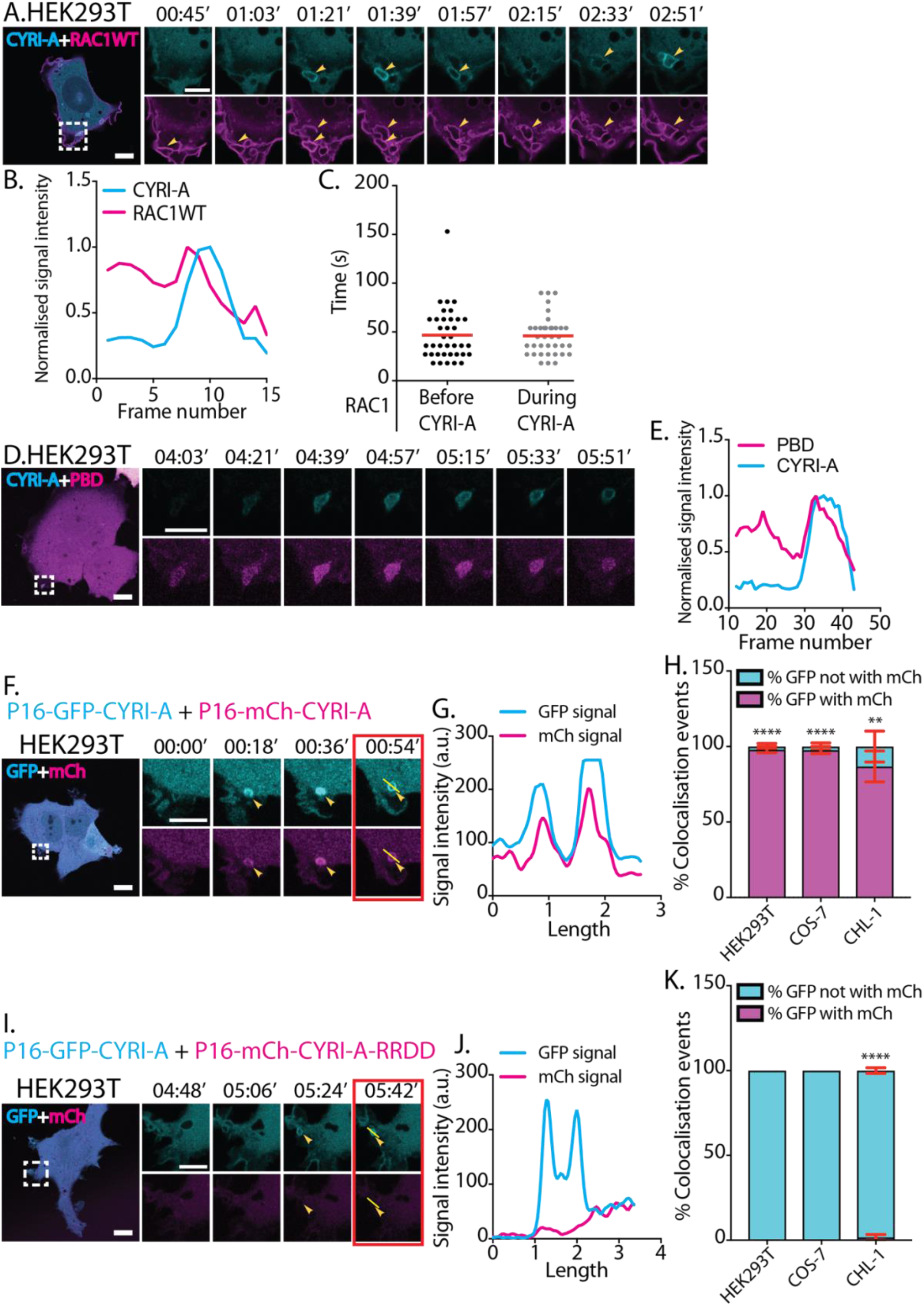
CYRI-A is recruited to macropinocytic cups by active RAC1. A-C. Time sequence images of HEK293T cell expressing P16-mCherry-CYRI-A and GFP-RAC1 WT. (B) shows the normalised signal intensity of the sequential event between RAC1 and CYRI-A. (C) Quantification of the lifetime of RAC1 signal on the macropinosomes (N=37 events in 4 cells). Scale bar = 10μm for full-sized image and 5μm for zooms. See also Video 11. D-E. Time sequence images of HEK293T cell expressing P16-GFP-CYRI-A and CFP-PBD. The graph shows the normalised signal intensities of CYRI-A and PBD over time. Scale bar = 10μm for full-sized image and 5μm for zooms. See also Video 12. F-K. Time sequence images of HEK293T cells expressing either WT or RRDD mutant of P16-mCherry-CYRI-A with the WT P16-GFP-CYRI-A (F, I). Line graphs show the colocalization of the signals between the two fluorescent constructs (G, J). Bar charts show the quantification of the percentage of colocalization events (N=5 cells) See also Videos 13-14 (H, K). Scale bar = 10μm for full-sized image and 5μm for zooms. Statistical analysis using two-tailed unpaired t-test. **p<0.01, ****p<0.0001.

### The recruitment of CYRI-A to macropinosomes is downstream of and dependent on PI3K signalling

PI-3 Kinase signalling plays an important role during micropinocytosis. PIP3 recruits RAC1 GEFs to macropinocytic cups to mediate RAC1 activation (Araki et al., 2007; Bloomfield and Kay, 2016; Bohdanowicz and Grinstein, 2013; Campa et al., 2015). We visualised phosphatidylinositol (3,4,5)-trisphosphate (PIP3) using two independent reporters PH-Grp1 (Kavran et al., 1998; Lai et al., 2013) and PH-Btk (Araki et al., 2007; Varnai et al., 2005) along with P16-GFP-CYRI-A in HEK293T cells. Remarkably, the PIP3 signal appeared for approximately 40 sec and peaked before the CYRI-A signal slowly appeared **(Fig. 7 A-F, Videos 15-16**). Similar observations were made in COS-7 cells, even though the PIP3 reporters appeared to coalesce on long tubules clumping together, which made spatiotemporally resolving them more difficult (**Fig. S3 G-L, Videos 32-34**). Our observations fit nicely with the consensus model where newly generated PIP3 recruits RAC1 GEFs, to activate RAC1 and drive actin polymerisation (Bloomfield and Kay, 2016; Buckley and King, 2017; Swanson and Watts, 1995) and our results suggest that the recruitment of CYRI-A can dampen down this signalling pathway. Blocking PI3K activity using 20μM LY294002 almost completely disrupts the recruitment of CYRI-A to the vesicles in COS-7 **(Fig. 7 G-H**, **Videos 17-18**). It is interesting to note that cells treated with LY294002 still retain membrane ruffles (Araki et al., 1996), but both diffused and localised recruitment of CYRI-A, is completely abolished. This fits in with a previous observation that LY294002 blocks the resolution of macropinosomes (Araki et al., 1996). Interestingly, when we label the plasma membrane in HEK293T cells using mScarlet-Lck (Chertkova et al., 2020), we observe that the plasma membrane invaginates for 50s before CYRI-A is recruited (**Fig. S3 M-N**). Just before the completion of the vesicles, the CYRI-A signal appears coincident with the budding off of the vesicles into the cytoplasm. This suggests that the CYRI-A signal is distinct from the very early step during the invagination process and perhaps also hints to its role in the budding off process of macropinocytic cups into vesicles. Overall, the recruitment of CYRI-A to macropinocytic cups is downstream of and strongly dependent on PI3K-PIP3 activity.

**Fig. 7.**
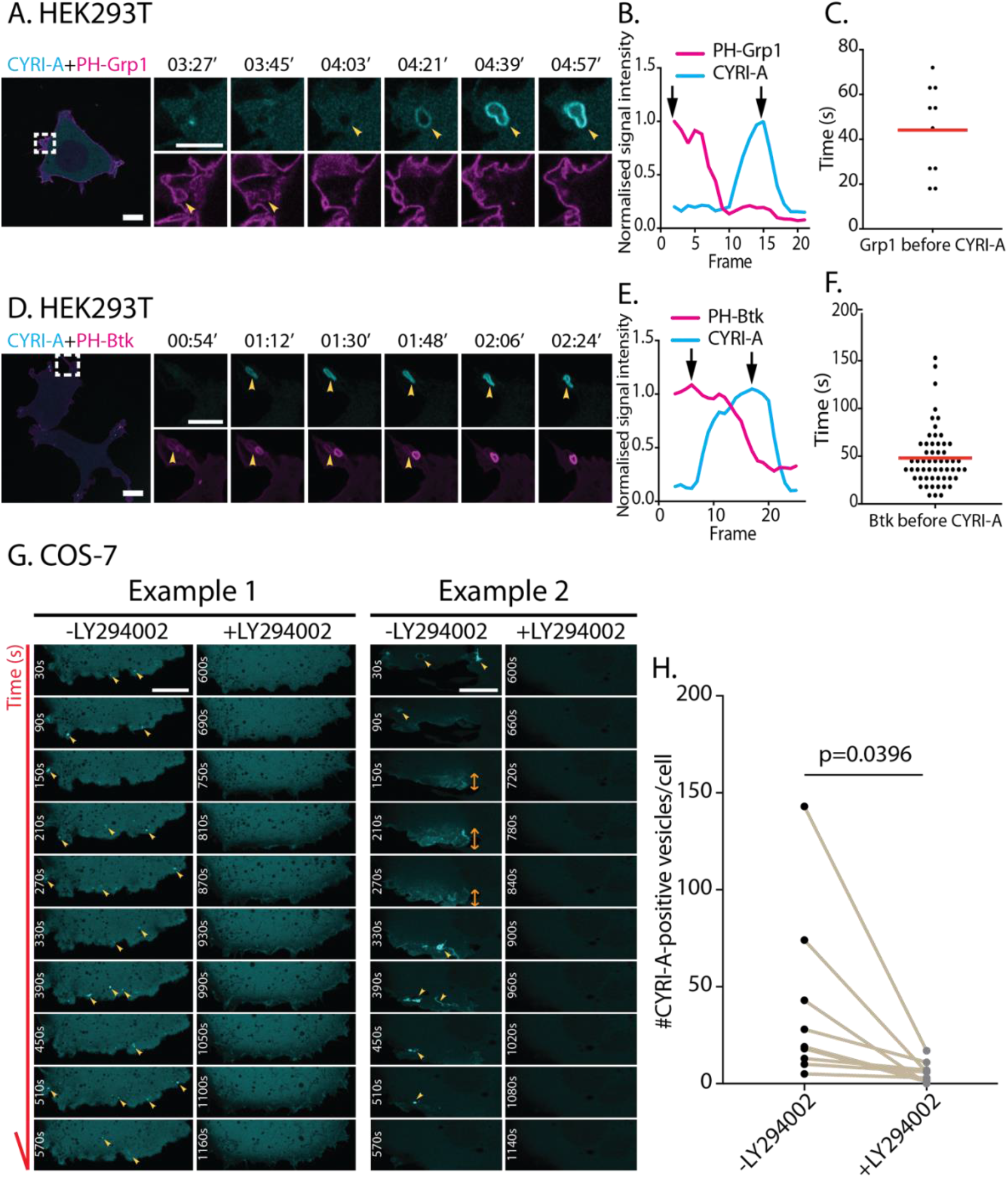
CYRI-A’s recruitment to macropinocytic cups is downstream of and dependent on PI3K signalling. A-F. HEK293T cells were co-transfected with mCherry-tagged CYRI-A and either GFP-PH-Grp1 or GFP-PH-Btk as specific markers for Phosphatidylinositol-3,4,5-phosphate (PIP3) (A, D). Line graphs show the sequential events between PIP3 reporters and CYRI-A (B, E). Black arrows indicate the peaks of each normalised signal. Scatter plots show the average lifetime of PIP3 reporter signal before CYRI-A is recruited to the macropinosomes (Grp1: N=9 events in 3 cells; Btk1: N=57 events in 6 cells). Red lines represent the average value. Scale bar = 10μm for full-sized images and 5μm for zooms. See Videos 15-16. G-H. Time sequence images showing COS-7 cells expressing GFP-tagged CYRI-A before and after the addition of 20μM of LY294002. Quantification shows a significant decrease in the number of vesicles formed over time (N=9 cells). Scale bar = 10μm. Statistical analysis using paired t-test. See also Videos 17-18.

### Integrin α5β1 is a major cargo of CYRI-A mediated macropinocytic uptake

One of the major cargoes of macropinocytic uptake is integrins (Gu et al., 2011; Caswell, 2009; De Franceschi, 2015). Furthermore, the broad lamellipodial phenotype of CYRI-A/B knockout cells (DBKO) suggested a possible enhancement of adhesion. We hence queried whether CYRI proteins could impact on the adhesion and integrin trafficking. Using the xCELLigence adhesion assay system based on electrical impedance, we found that CYRI DBKO A-673 cells spread around 2-fold faster than single knockouts or controls **(Fig. 8 A-B**). Flow cytometry analysis revealed an almost 50% increase in the surface expression of integrin α5 and β1 in the two independent DBKO A-673 cell lines compared to the control pLKO **(Fig. 8 CD**). Western blotting shows a minor but consistent increase of the total level of integrin α5 and β1 but qRT-PCR analysis shows no obvious change in their mRNA levels **(Fig. 8 E-F**). We detected a similar increase for the active integrin α5 using the SNAKA51 antibody with flow cytometry, but no change for the level of the membrane metalloprotease MT1-MMP (**Fig. S4 A-B**). Immunofluorescence analysis revealed a significant increase in both the area of adhesion sites as well as their number per cell in the DBKO cells compared to the controls **(Fig. 8 G-H**, Fig. S4 C-E). Overexpression of CYRI-A in the DBKO cells rescued these phenotypes **(Fig. 8 I-J**). Overall, we find that loss of both CYRI isoforms in A-673 cells leads to the enhanced display of surface integrins, suggesting a defect in their intracellular trafficking.

**Fig. 8.**
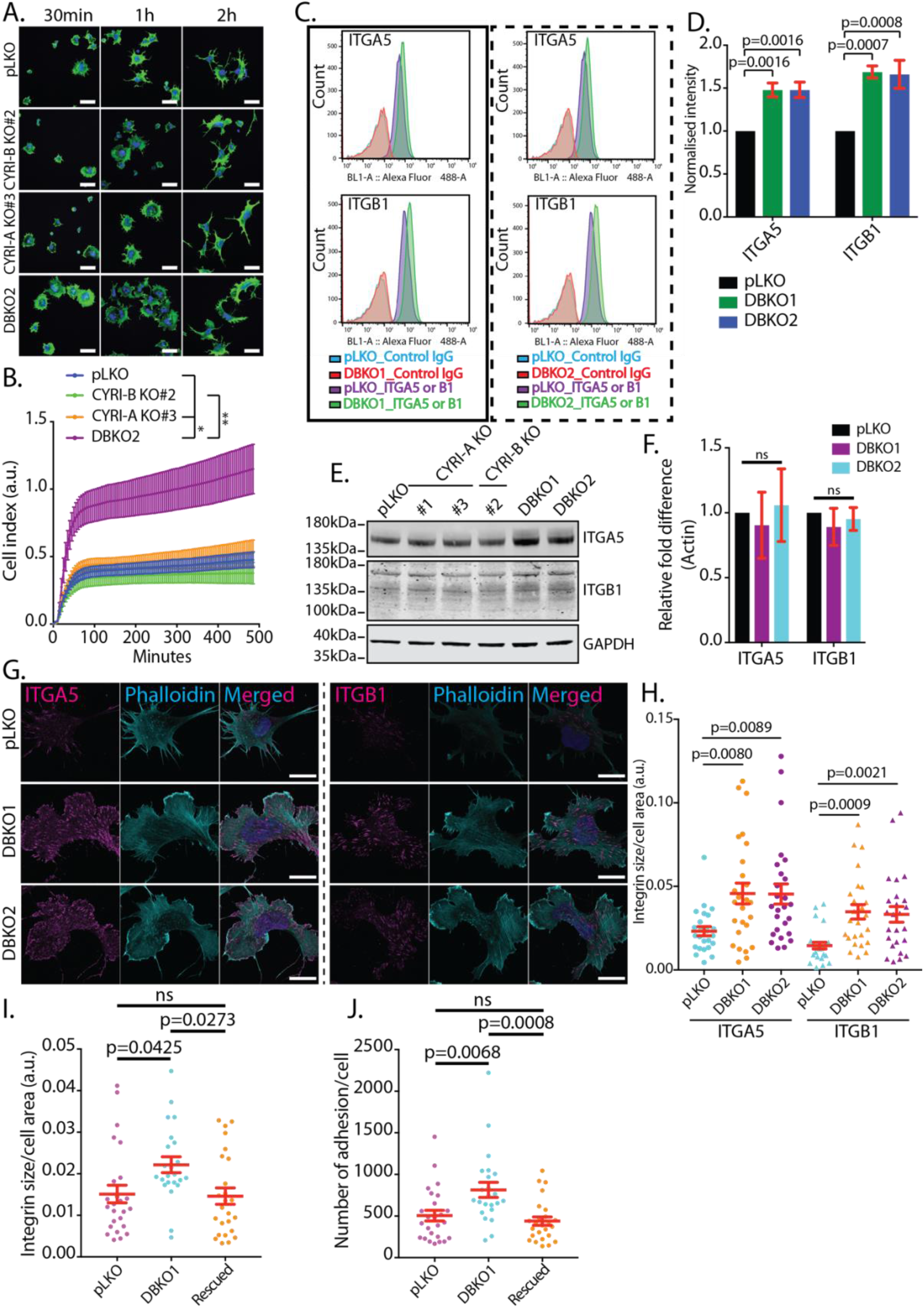
CYRI-A and CYRI-B cooperatively regulate matrix attachment and surface display of integrin α5β1. A-B. Representative immunofluorescence images of A-673 comparing the spreading propensity between the control pLKO and the DBKO cells on fibronectin (A). Scale bar = 40μm. Graph shows a time course of cell-matrix attachment for control pLKO (blue), the single knockout (green and orange) and the DBKO cells. C-D. Flow cytometry analysis (C) and quantification (D) of the surface expression of integrin α5β1 in A-673 cells between control pLKO and the DBKO cells (DBKO1 and DBKO2). E-F. Representative western blot of the total level of Integrin α5β1 in DBKO cells compared to control pLKO or single knockout cells (E). qPCR analysis of the gene expression of integrin α5β1 between pLKO cells and the DBKO cells (F). G-H. Immunofluorescence images of the surface level of integrin α5β1 between control pLKO and DBKO cells (#1 and #2) (G). Scale bar = 10μm. The quantification is shown in H. Data from at least 10 cells per experiment. I-J. Effect of CYRI-A expression in DBKO cells on area and number of integrin adhesion sites. Data from at least 10 cells per experiment. Data from 3 independent experiments. Mean ± SEM. Statistical analysis using ANOVA with Tukey’s multiple comparison test. *p<0.05, **p<0.01.

### Integrin α5 and β1 are recruited to CYRI-positive cups and vesicles

We next asked whether integrins colocalize with CYRI proteins at these structures. We first co-expressed a mApple-tagged integrin α5 construct along with the P16-GFP-CYRI-A or P17-GFP-CYRI-B in COS-7 cells **(Fig. 9 A-B, Videos 19-20**) or HEK293T cells (**Fig. S4 F, Video 35**). Integrin α5 decorated many intracellular vesicles, including CYRI-A-positive macropinocytic cups formed near the edge of the cells **(Fig. 9 A**). CYRI-B decorated endocytic tubules emanating from the tip of filopodia-liked protrusions or from the side of the cell **(Fig. 9 B**). Endogenous integrins also colocalized with CYRI-A and CYRI-B-positive macropinocytic cup-like structures in COS-7 **(Fig. 9 C-H**) as well as in the DBKO A-673 cells (**Fig. S4 G-J**).

**Fig. 9.**
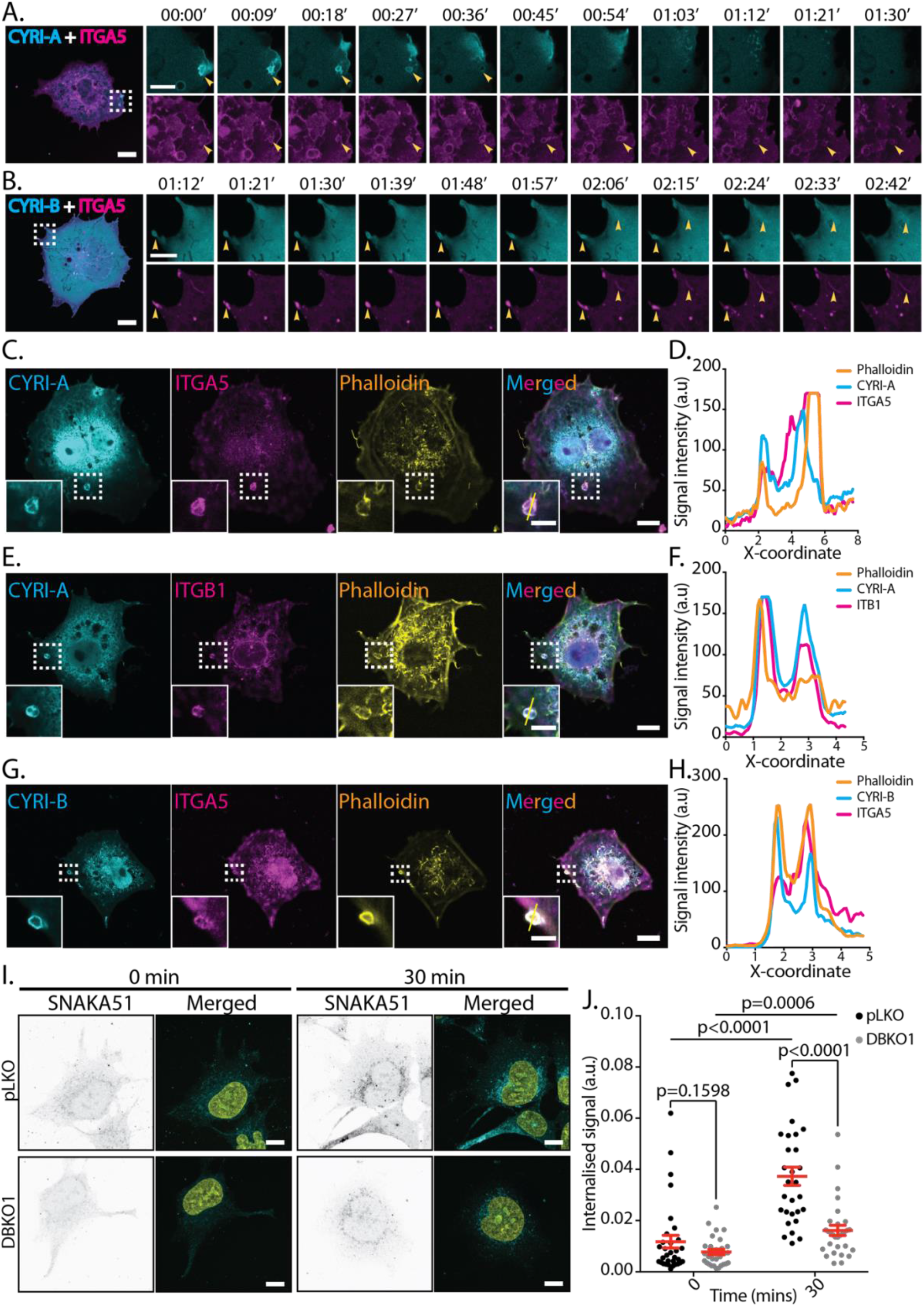
CYRI-A and CYRI-B cooperatively regulate the internalisation of integrin α5β1. A-B. Time sequence images of live COS-7 cells expressing either P16-GFP-CYRI-A or P17-GFP-CYRI-B and mApple-tagged integrin α5. Scale bar = 10μm for full-sized image and 5μm for zooms. C-H. Colocalization analysis of CYRI-A or CYRI-B (cyan) with endogenous integrin α5 or β1 (magenta) and actin (orange) in COS-7 cells. Right panels (D, F, H) are intensity graph showing the colocalization of all three normalised signals at the vesicles. Scale bar = 10μm for full-sized image and 5μm for zooms. I-J. Internalisation assay of A-673 cells for active integrin α5 (SNAKA51, cyan) after 0- or 30-minute incubation at 37°C, (nuclei, DAPI, yellow). Left panels are representative images showing the internalised signal of active integrin α5 (SNAKA51, black signal). The right panel is the quantification of the area of the internalised signals. Data from 3 independent experiments of at least 10 cells per experiment. Statistical analysis using two-tailed unpaired t-test. Scale bar = 10μm.

Increased surface presentation of integrin α5 and β1 suggested a potential defect in internalisation, perhaps due to the requirement for CYRI proteins in the resolution of macropinocytic structures. To address this, we performed an internalisation assay of the active integrin α5. After 30min of induction, we observed a significant increase in the internalised SNAKA51 signal, particularly in the control cells (**Fig. 9 I-J**). Also, the control pLKO cells have a much stronger intracellular signal compared to the DBKO cells, suggesting that a higher level of active integrins were internalised. Overall, our data support a novel function of CYRIs in cooperatively regulating the trafficking of integrin α5β1 through the enhancement of actin dynamics during macropinocytosis to allow the efficient internalisation of active surface integrins.

### CYRI depletion promotes invasiveness and resistance to anoikis due to enhanced active surface integrins

Increased level of α5 and β1 integrins have been associated with increased invasiveness and worse outcome for multiple types of cancer (Bianchi-Smiraglia et al., 2013; Clark, 2005; Mierke, 2011; Nam et al., 2010; Paul et al., 2015). Building on these observations and our migration data **(Fig. 3 D-H**, **Fig. S1 F-G**), we tested the invasive capacity of the CYRI DBKO A-673 cells using an organotypic assay where cells invade into native collagen plugs (Timpson et al., 2011). Loss of CYRI proteins significantly increased invasion **(Fig. 10 A-B**). The invasion index of the DBKO cells (~10%) is >3-fold above controls (~3%). Blocking the activity of the integrin α5β1 using the blocking antibody IIA1 dramatically reduced the invasion of the DBKO cells but not the control cells into collagen-Matrigel-fibronectin (CMF) plugs in inverted invasion assays **(Fig. 10 C-D**). Individually inhibiting each subunit with specific blocking antibodies also results in the same reduction (**Fig. S5 A-B**). On a 2D surface, pre-treating cells with IIA1 blocking antibody significantly reduces their ability to adhere and spread on fibronectin matrix (**Fig. S5 C-D**). This suggests that in CYRI DBKO cells, integrins play a crucial role in regulating their shape and adhesion. Treatment with this antibody, however, only affects the migration ability of DBKO cells but not the control pLKO in a 2D random migration assay (**Fig. S5 E**), which is in agreement with our invasion data **(Fig. 10 C-D**). Integrins also provide cancer cells with increased resistance to anoikis, a programmed cell death process triggered by the lack of adhesions, through triggering FAK activation on the endosomal membrane (Alanko et al., 2015). Culturing the control and two DBKO cell lines in the low-attachment condition of agarose shows that indeed the DBKO cells form larger colonies **(Fig. 10 E-F**). Overall, our data provide a mechanism linking the function of CYRIs in macropinocytosis to the invasive capacity of cancer cells through modulating integrin trafficking and signalling (**Fig. S5 F**).

**Fig. 10.**
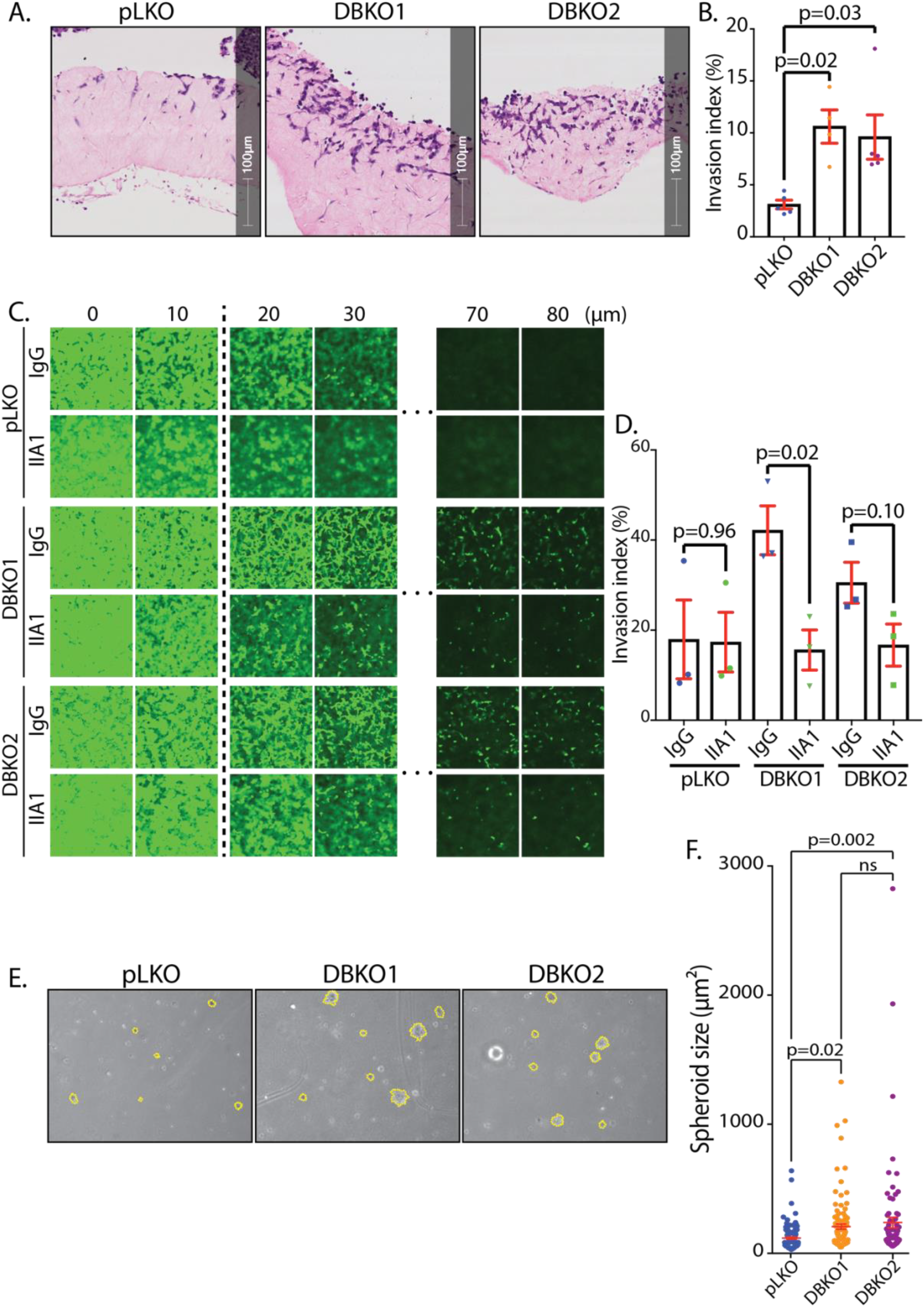
CYRI-depleted cells show enhanced invasion and anchorage independent growth due to increased active surface integrins. A-B. Organotypic assay of control pLKO versus CYRI DBKO A-673 cells stained with H&E. The graph shows the quantification of the invasion index (the area invaded by cells normalised to the total area of the plug). Data are from at least 3 independent experiments. Mean ± SEM. Statistical analysis using ANOVA with Tukey’s multiple comparison test. C-D. Inverted invasion assay comparing the invasion capacity between the control pLKO and the CYRI DBKO A-673 cells in the presence or absence of the integrin α5β1 - blocking antibody IIA1. The graph shows the quantification of the invasion index (the area covered by cells beyond 10μm into the plug). Data are from at least 3 independent experiments. Mean ± SEM. Statistical analysis using two-tailed unpaired t-test. E-F. Phase-contrast images of the soft agar assay comparing the anchorage-independent growth of the control pLKO and the DBKO cells. The size of the spheroids is quantified in F. Data are pooled from 3 independent experiments. Mean ± SEM. Statistical analysis using ANOVA with Tukey’s multiple comparison test.

## Discussion

CYRI proteins show activity as negative regulators of RAC1-mediated activation of the Scar/WAVE complex, but nothing is known about their dynamic relationship with RAC1 and actin at the plasma membrane. Furthermore, the cellular functions of CYRI-A have never been specifically described. CYRI-A, encoded by the *FAM49A* gene, has previously been linked to non-syndromic oral clefts and craniofacial abnormalities (Azevedo et al., 2020; Chen et al., 2018; Leslie et al., 2016), suggesting an important role in developmental and perhaps morphogenetic processes, but its cellular and molecular function has not been studied. Here, we model CYRI-A into the recently solved structure of CYRI-B (Kaplan et al., 2020; Yelland et al., 2020) and we conclude that CYRI-A likely adopts a similar structure to CYRI-B but may also have important differences. We show biochemically that CYRI-A interacts with active RAC1 using similar amino acid residues, but with a higher affinity than CYRI-B. CYRI-A expression can rescue the effects of CYRI-B deletion on lamellipodia and spreading, reflecting that they likely have overlapping functions. Depletion of both CYRI-A and CYRI-B together has a stronger effect on lamellipodia and cell migration than depletion of either protein independently. We thus place CYRI-A as a paralog of CYRI-B, with broadly similar behaviour biochemically and in cells. Here, we examine for the first time, the dynamics of CYRI proteins in live cells and reveal an unexpected dynamic role of CYRI-A in macropinocytosis, where it drives resolution of the macropinocytic cups, with implications for integrin trafficking, cell migration and invasion.

The interplay between signalling and cytoskeletal dynamics that drives macropinocytosis involves PI3K activation and PIP3 signalling leading to the activation of RAC1 (likely via recruitment of a RAC1 GEF) and thus the Scar/WAVE complex. The resulting actin assembly drives the formation of membrane ruffles and macropinosomes (Veltman et al., 2016). We observed a dynamic recruitment of CYRI-A at macropinocytic cups, persisting until after engulfment. Using markers of the various stages of macropinocytosis, we place CYRI-A downstream of PI3K, RAC1 activation and the initiation of actin assembly at the nascent macropinocytic cup (see Model, **Fig. S5F**). By contrast, CYRI-B appeared to be more evenly distributed at the plasma membrane and on internal tubular and vesicular structures. So, while CYRI-A and CYRI-B can compensate for each other in aspects of cell motility, they appear to be differently regulated.

Both the activation and deactivation of RAC1 are required for the completion of macropinocytosis. For example, when RAC1 activation is sustained using photoactivation, the macropinocytic cups were arrested until RAC1 activation was switched off (Fujii et al., 2013). In macrophages, RAC1 GAPs are frequently paired with GEFs (Denk-Lobnig and Martin, 2019) and are needed for the completion of phagocytosis of large particles (Schlam et al., 2015), presumably to shut down RAC1 signalling and to allow actin disassembly. However, a specific inhibitor for RAC1 at the macropinocytic cup has never been discovered. Here, we show that CYRI-A rapidly accumulates on macropinocytic cups harbouring activated RAC1, where it antagonises RAC1 activity toward Scar/WAVE, followed by a decrease in the actin signal. Importantly, CYRI-A and actin form a feedback loop, where blocking actin dynamics using Latrunculin A or Cytochalasin D shortened CYRI-A’s lifetime on macropinosoomes. Conversely, CYRI-A decreases the lifetime of accumulated Factin on macropinosomes, indicating reciprocal feedback. Interestingly, a proteomic analysis of macropinosomes done in *Dictyostelium* also implicated CYRI (then only known as Fam49) (Journet et al., 2012), suggesting that this function is likely to be conserved. How CYRI-A interplays with RAC1 GEF and GAP proteins is unknown, but our evidence places CYRI-A as a specific locally-acting disruptor of RAC1 – Scar/WAVE signalling.

Double knockdown of CYRI-A and CYRI-B gave a robust enhancement of cell spreading and concomitant increase in surface α5β1 integrin. Integrins are internalised by multiple different routes, including macropinocytosis (Gu, et al., 2011) and the CLIC-GEEG pathway (Moreno-Layseca et al., 2020), where they can be rapidly recycled back to the surface or targeted for degradation in lysosomes (Yu et al., 2012; Zech et al., 2011). It is unclear whether certain routes of uptake such as clathrin-mediated endocytosis or macropinocytosis, might favour degradation versus recycling, but we find that by depleting CYRIs, we slow down the internalisation of active integrin as detected with the SNAKA51 antibody (**Fig. 9**). The SNAKA51 antibody preferentially recognises active integrin α5β1 at fibrillar adhesions that are bound to fibronectin (Clark et al., 2005). Active ligand-bound α5β1 integrin is also internalised via an Arf4-Scar/WAVE-dependent manner for trafficking to lysosomes (Rainero et al., 2015). Integrin engagement and recycling are important regulators of cancer cell invasive migration (Caswell et al., 2008; Dozynkiewicz et al., 2012; Paul et al., 2015; Yu et al., 2012). Loss of CYRI-A and -B promoted cancer cell invasion, presumably due to excess active integrins on the cell surface and perhaps higher overall levels of RAC1 activation and engagement of the Scar/WAVE complex at the cell’s leading edges.

We thus find that while CYRI-A has the ability to rescue the cytoskeletal defects of CYRI-B knockout cells, it shows a striking and highly dynamic localisation to macropinocytic cups and colocalizes with integrins during their internalisation via macropinocytosis. We show that CYRI proteins regulate integrin homeostasis and loss of CYRI-A and B in a double knockout mutant leads to excessive surface integrin and an increase in invasion. Our results emphasise that activation of the Scar/WAVE complex by RAC1 not only affects cell migration and/or invasion via altering actin dynamics at the leading edge of cells, but that this pathway, regulated by CYRI proteins, is a major regulator of surface integrin levels. This could have important implications for cancers where RAC1 pathways are disrupted or enhanced, such as in melanomas where RAC1 activating mutations are considered drivers (Cannon et al., 2020).

## Materials and methods

### Cell lines generation, maintenance and growth conditions

COS-7, CHL-1 and HEK293T cells were grown in DMEM (Gibco, #21969-035) supplemented with 10% fetal bovine serum (Gibco, #10270-106) and 1X glutamine (Gibco, #25030-032) and 1X Pen/Strep (Life Technologies, #15140122). A-673 cells were a kind gift from Dr Susan Burchill, Leeds, UK and were grown in the same media but with 0.5X glutamine. Single CYRI CRISPR A-673 was generated using Lentiviral CRISPR vector (hSpCas9-2A-Puro, Addgene #62988) and selected using 1μg/ml Puromycin (Invivogen, #ant-pr-1) for 1 week. Double CYRI CRISPR A-673 was generated using the combination of the hSpCas9-2A-Puro and a modified version containing a Blasticidin resistant gene (a kind gift from Dr Stephen Tait, CRUK Beatson Institute, Glasgow, UK). Double knockout cells were selected by using a combination of 1μg/ml Puromycin and 6μg/ml Blasticidin (Invivogen, #ant-bl-10p) for 1 week.

For maintenance, cells were kept in the presence of all antibiotics to prevent incomplete selection. For passaging, cells were washed with PBS and 500μl of 0.25% trypsin (Gibco, #15090046) was added and incubated at 37°C for 2min. Trypsin was blocked by adding 5ml of medium containing 10%-serum and cells were split in a 1:10 ratio 2 times a week. For experimental testing, cells were grown in antibiotic-free media. Cells were grown at 37°C with 5% CO2 incubator.

For all experiments with multiple repeats, cells of different passages were considered biological replicates. Cells of different origins (HEK293T, COS-7, CHL-1) were used to confirm for the reproducibility of our observations in live-cell imaging experiments. All cells except A-673 were from ATCC and were authenticated using STR profiling. All cells were tested for mycoplasma and were confirmed to be negative.

### sgRNA and siRNA

sgRNAs for CRISPR were designed using the Zhang lab website (https://zlab.bio/guide-design-resources) and were then purchased through ThermoFisher Scientific.

h49A-sgRNA2.1

5′-CACCGCCTGAAGGACGGCGCTGATC-3′

h49A-sgRNA2.3

5′-CACCGCTGCAGGCTTACAAAGGCGC-3′

h49B-sgRNA4.1

5′-CACCGCGAGTATGGCGTACTAGTCA-3′

siRNAs are purchased from Qiagen.

Hs_FAM49A_5 FlexiTube siRNA (#SI03150210)

Hs_FAM49A_9 FlexiTube siRNA (#SI05122656)

Hs_FAM49B_7 FlexiTube siRNA (#SI04359369)

AllStars Neg. Control siRNA (20nmol) (#0001027281)

For CRISPR gene deletion, lentivirus containing the CRISPR-Cas9 construct was generated using HEK293T cells. In brief, 1.5×10^6^ HEK293T cells were plated on a 10cm Petri dish. Cells were transfected with 10μg of CRISPR construct containing sgRNA of the gene of interest, 7.5μg of pSPAX2 and 4μg of pVSVG packaging plasmid using standard calcium precipitation protocol (Sigma) and grown in 20% serum-containing medium for 24h. Conditioned media were filtered through a 0.45μm filter and mixed with polybrene before being added to the targeting cells. The infection process was repeated a second time before antibiotics were added during 1 week for selection.

For siRNA gene silencing, for 1 reaction, 20nmol of siRNA was mixed with 7μl of Lullaby (Oz Biosciences, #LL71000) and incubated in serum-free medium in a total volume of 200μl for 20min before adding to 3×10^5^ cells growing in 1800μl of medium. The process was repeated again after 48h and cells were split if becoming too confluent. 24h after the 2^nd^ treatment, cells were used for further experiments.

### Lipofectamine plasmid transfection

For 1 reaction, 2μg of DNA plasmid was mixed with 5μl of Lipofectamine 2000 (ThermoFisher Scientifics, #11668019) in serum-free medium to a total volume of 200μl and incubated at room temperature for 5min before adding to 3×10^5^ cells in a 6-well plate.

### Cell fixation and Immunofluorescence

Glass coverslips were treated in concentrated HNO3 (nitric acid) for 30min before washing with a copious amount of water. Coverslips were then coated with 1mg/ml of fibronectin (Sigma-Aldrich, #F1141) for 2h and washed with PBS. Cells were seeded on coverslips and left to settle for 4h before being fixed in 4% paraformaldehyde (PFA). Cells were permeabilised with buffer containing 20mM glycine, 0.05% Triton X-100 in PBS for 5min before being incubated with primary and secondary antibodies. Cells were then mounted on to a glass slide using ProLong™ Diamond Antifade Mountant (Invitrogen, #P36961).

### CYRI-B CRISPR COS-7 cell CYRI-A rescue experiments

CYRI-B CRISPR COS-7 cells were transfected with a control GFP-FLAG or CYRI-A-FLAG construct for 24h before cells were seeded on acid-treated fibronectin-coated coverslips for 4h. Cells were fixed and stained using the immunofluorescent protocol described above. The average Arp2/3 signals at the cell periphery of the transfected cells were measured using the freehand selections tool with the width of 1 pixel.

### Integrin adhesion immunofluorescent quantification

The number and size of integrin adhesion sites were extracted using auto threshold function in ImageJ. The number of integrin adhesion sites was the absolute number per cell. The size was normalised to the size of the selected cell.

### Image-based Integrin internalisation assay

The assay was based on the internalisation assay described in (Pietila et al., 2019) with a slight modification. In brief, A-673 cells were grown on fibronectin-coated coverslips overnight to allow the cells to fully adhere. The next day, cells were washed with ice-cold PBS and incubated with SNAKA51 antibody in ice-cold Hank’s Balanced Salt Solution (HBSS) buffer (8g NaCl, 0.4g KCl, 0.14g CaCl2, 0.1g MgSO4.7H2O, 0.1g MgCl2.6H2O, 0.06g Na2HPO4.2H2O, 0.06g KH2PO4, 1.0g glucose, 0.35g NaHCO3, H2O to 1000ml) for 1h on ice and covered in aluminium foil. Internalisation was induced by adding 1ml of 37°C full serum growth medium and quickly transferred to an incubator for 30min. For the control as 0min, the unbound antibody was washed with ice-cold PBS before the bound antibody was stripped using acid wash (0.2M acetic acid, 0.5M NaCl, pH 2.5) for 1.5min. After 30min, antibodies on the internalised cells were stripped and both conditions were fixed with 4% PFA. Cells were then subjected to our standard immunofluorescence protocol as described above. For image acquisition, a z-stack of 0.18μm per slice was taken with the AiryScan Zeiss880 using the Plan-Apochromat 63x/1.4 oil DIC M27 objective lens. A maximum intensity projection was generated using the 3D Project function from ImageJ. The pipeline for semi-automated quantification was set up. Auto threshold function in ImageJ was used to identify internalized signals. Enhanced contrast images were used to define cell area. The level of internalization was quantified by measuring the area of internalized signal and normalized to the area of the selected cell.

### Macropinocytosis assay

Cells were transfected with siRNA targeting both *CYRI-A* and *CYRI-B* for 48h and 24h before seeding onto fibronectin-coated coverslips overnight. The next day, cells were washed three times with ice-cold PBS before being incubated with 0.2mg/ml dextran 70kDa 488 (ThermoFisher Scientific, #D1822) for 15 or 30min in serum-free medium at 37°C. Cells were then washed five times in ice-cold PBS and fixed for 15min using 4% PFA before staining with DAPI for 30min. Ten random fields of view were taken and analysed using semi-automated ImageJ macro. The auto threshold function in ImageJ was used to measure the internalised dextran signal. Enhanced contrast images were used to define the area of cell clusters. Macropinocytic index was calculated by normalising the Internalised dextran signal to the total area of the cell cluster for each field of view (Commisso et al., 2014).

### Western blotting

Cells were grown to 80% confluence in a 6-well plate and lysed with RIPA buffer (150mM NaCl, 10mM Tris-HCl pH7.5, 1mM EDTA, 1% Triton X-100, 0.1%SDS) supplemented with protease and phosphatase inhibitor cocktails (ThermoFisher Scientific, #78438, #78427). Lysates were spun at maximum speed for 10mins before protein level was measured using Precision Red (Cytoskeleton, Inc., #ADV02). Samples were run using precast 4-12% NuPAGE Bis-Tris Acrylamide Gels (ThermoFisher Scientific, #NP0321) and transferred to a 0.45μm nitrocellulose membrane (GE Healthcare, #10600002). Membranes were blocked in 5% BSA in TBS-T for 30min before incubated with specific primary antibodies and the corresponding secondary antibody (Invitrogen, #A21206 and #A10038; ThermoFisher Scientific, #SA5-35521 and #SA5-35571) before being analysed using the Image Studio Lite (LiCor system).

### GST-Trap Pulldown

For small scale GST-tagged protein purification, BL21 competent *E. coli* was grown until OD 0.4 and induced with 200μM IPTG overnight at room temperature. Cells were pelleted and lysed using sonication in buffer containing 50mM Tris 7.5, 50mM NaCl, 5mM MgCl2, 250μM DTT. The lysates were collected by ultracentrifugation at 20,000rpm for 20min. The lysates were added directly on glutathione sepharose 4B (GE Healthcare, #17075601) for 2h. COS-7 cells were transfected with GFP-tagged constructs overnight and lysed the next day using lysis buffer containing 100mM

NaCl, 25mM Tris-HCl pH7.5, 5mM MgCl2 supplemented with 0.5% NP-40 and protease and phosphatase inhibitors. Lysates were cleared by centrifugation and incubated with GST beads for 2h before subjected to western blotting as described.

### Recombinant protein purification and surface plasmon resonance (SPR)

In brief, BL21 competent *E. coli* containing MBP-tagged or GST-tagged protein were grown to OD 0.4 and induced overnight with 200μM IPTG at room temperature. The next day, cells were concentrated and lysed using the AKTA machines in lysis buffer (20mM Tris pH 8.0, 300mM NaCl, 5mM MgCl2 and 2mM beta-mercaptoethanol (BME)) supplemented with protease inhibitors. Cell lysates were cleared using ultracentrifugation at 20,000rpm for 45min. The lysates were passed through an affinity column either a MBPTrap HP (Sigma, # GE28-9187-79) or a GSTrap 4B (Sigma, # GE28-4017-47) (buffer contains 20mM Tris, 150mM NaCl, 5mM MgCl2 and 2mM DTT) before being eluded with either 10mM maltose or glutathione. The eluded fractions were then subjected to a size exclusion chromatography using an (HiLoad16/600 Superdrex 200pg). GST tags were digested overnight with thrombin and were then separated out using affinity chromatography.

For SPR: MBP-tagged proteins were immobilised on Sensor chip CM5 (GE Healthcare, #BR100012) and purified untagged RAC1 Q61L was titrated at different concentrations and the response signals were recorded and fitted into a 1:1 binding model. The dissociation constant Kd was estimated from the fitted curve.

### 2D and 3D CDM migration assay

6-well plates were coated with 1mg/ml of fibronectin for 2h before 20,000 cells were seeded and left for 4h to settle at 37°C. Cells were imaged using the Nikon TE2000 microscope equipped with a PlanFluor 10x/0.30 objective every 15min for 20h. Individual cells were tracked overtime manually using MTrack ImageJ plug-in and the velocity was calculated using Chemotaxis tool ImageJ plug-in.

Cell-derived matrix (CDM) was generated using Telomerase Immortalised Fetal Foreskin (TIFF) cells and kept in PBS at 4°C until use (Cukierman et al., 2001). The same seeding, imaging and analysis protocol was used as for the 2D migration assay.

### Wound-healing assay

A 96-well plate (Falcon, #353072) was coated with 1mg/ml fibronectin for 2h. Cells were seeded to form a confluent monolayer (70,000 cells for A-673) and left to settle for 4h. The wound was made using the Incucyte wound maker. Cells were stained with IncuCyte Nuclight Rapid Red reagent (#4717) for total cells and IncuCyte Sytox Green reagent (#4633) for dead cells. Cells were imaged every hour for 48h using the IncuCyte system. Analysis of wound closure was done on the Zoom software (IncuCyte Biosciences, Sartorius).

### Proliferation assay

1×10^4^ cells were seeded per well in a fibronectin-coated 96-well plate (Falcon, #353072). Cells were stained with IncuCyte Nuclight Rapid Red reagent (#4717) for total cells and IncuCyte Sytox Green reagent (#4633) for dead cells and were imaged for 28h. To calculate the growth curve, the total number of alive cells were calculated by subtracting the number of Sytox-positive cells from the number of Nuclight Red-positive cells. This was then normalised to the number of alive cells from time 0. The slope is calculated using a non-linear regression model from the software Prism7.

### xCELLigence spreading assay

xCELLigence E-plate 16 (Acea, #05469830001) was coated with fibronectin for 1h. Cells were trypsinised as described earlier, counted and 10,000 cells were seeded on the plate. Immediately the plate was moved to an Acea RTCA DP xCELLigence device and cell spreading was measured based on electrical impedance every 5min for 8h.

### Live-cell super-resolution fluorescence microscopy

For all live imaging experiments, cells were seeded on fibronectin-coated 3.5cm dish for 4h prior to imaging. During the imaging process, cells were kept in a 37°C with 5% CO2 chamber. Imaging was performed with a Zeiss 880 Laser Scanning Microscope with Airyscan with a Plan-Apochromat 63x/1.4 oil DIC M27 objective lens and deconvoluted using a Zen Black Zeiss880 deconvolution software (ZEN 2.3 SP1 FP3 v14.0.17.29).

For actin depolymerisation experiment, at least 5 cells were imaged before Latrunculin A (Sigma-Aldrich, #L5163) or Cytochalasin D (Sigma-Aldrich, #C2618) were added at 1 μM. Then, at least 5 random cells were imaged after at least 5 min of drug treatment. Images were taken every 9s.

For PI3K inhibition experiment, 9 random cells were imaged every 30 seconds for 20 frames before LY294002 was added at 20μM and the imaging process continued for another 20 frames.

For live-cell dextran incorporation assay, cells were first imaged every 9 seconds before dextran (Texas Red™ 70,000 MW Lysine Fixable, Invitrogen, #D1864) was added at 50μg/ml mid-imaging.

For all line graphs representing the signal intensity overtime, each line was normalised to the highest signal value.

To calculate the percentage of colocalization events, the number of vesicles colocalize or not colocalize between two signals were counted.

% colocalization events = (#colocalising events/#non-colocalising events)*100.

### Flow cytometry analysis

10×10^6^ cells were grown in a 15cm Petri dish overnight. The next day, cells were trypsinised, washed in PBS and filtered through a 40μm filter. All conditions were stained with Zombie Red reagent (#423110) before being fixed in 4% PFA for 15min. Cells were blocked in PBS supplemented with 10% serum for 30min and then incubated with the corresponding primary antibody for 15min at room temperature. If the primary antibodies were not conjugated with Alexa 488, cells were then incubated with the secondary antibody for 15min at room temperature before analysing using Attune NxT system (ThermoFisher Scientific). For negative control, cells were incubated with anti-human mouse IgG (BioLegend, #400132). For all conditions, dual colours of the far-red and 488 flow cytometry was used to minimise spectrum overlapping. Data were imported and analysed using FlowJo software (Becton Dickinson & Company, v. 10.6.1).

### Quantitative real-time PCR (qRT-PCR)

Cells were grown to 80% confluence in a 6-well plate before RNA was extracted using standard RNA extraction protocol (RNAeasy Mini Kit, Qiagen, #74104). cDNA synthesis was performed by mixing 1μg of RNA with qScript^®^ cDNA Synthesis Kit (Quantabio, #95047-100) and the reaction condition is as follows: 5min at 25°C, 30min at 42°C, 5min at 85°C using a thermocycler. qPCR was performed with the DyNAmo HS SYBR Green qPCR kit (ThermoFisher Scientific, #F410L) according to the manufacturer protocol and was done using the QuantStudio Real-Time PCR system. All primers were purchased from Qiagen website: Hs_ITGA5_1_SG QuantiTect Primer Assay (#QT00080871), Hs_ITGB1_1_SG QuantiTect Primer Assay (#QT00068124), Hs_ACT_2_SG QuantiTect Primer Assay (#QT01153551).

### Organotypic assay

The organotypic assay was performed based on the published protocol by Dr Paul Timpson (Timpson et al., 2011). In brief, collagen was extracted from rat tail and kept in 0.5mM acetic acid buffer at 2mg/ml concentration. Collagen plugs were made by mixing TIF cells with 25ml of collagen and the pH was adjusted using NaOH to 7.2. Collagen plugs were allowed to contract for 8 days before being transferred to a 24-well plate. A-673 cells were prepared as usual without any trypsin present in the final solution and 3×10^4^ were seeded directly on top of each plug. Media were changed every 2 days for 5 days before the plugs were transferred onto a metal grid. A liquid-air interface was set up with the media acting as a chemoattractant for the invasion. Media were changed every 2 days for 14 days before the plugs were being collected and processed for H&E histology staining. Images were quantified using HALO software (PerkinElmer). Invasion index was quantified by measuring the invading area and normalised it to the size of the entire tissue section.

### Inverted invasion assay

The plug was made by mixing Collagen I with Matrigel (Beckton Dickinson, #354234), fibronectin (CMF) in ice-cold PBS to the final concentration of 4, 4, 1 mg/ml respectively. 100μl of the mixture was added to each Transwell (Fisher Scientific, TKT-525-110X, C3422) with 8μm pore size and left to solidify at 37°C. 5×10^4^ A-673 cells were seeded on to the underside of the Transwell and left to adhere for 4h at 37°C. After this, the Transwell was inverted and dipped into 500μl of 10% serum-containing medium. The top chamber was filled with 100μl of serum-free medium. Inhibiting antibodies were added to the top chamber at a concentration of 5μg/ml. The cells were left to invade for 5 days before staining with 4μM Calcein AM (Invitrogen, #C1430). Z-stack images were captured every 10μm using the Olympus FV1000 confocal microscope with the UplanSApo 20x/0.74 objective. Invasion index was calculated by measuring the area of invading cells beyond 10μm in the plug and then normalised to the total area of invading cells. Area measurement was performed auto Threshold function in ImageJ.

### Low-attachment assay

Agarose was dissolved in PBS at 4% and autoclaved. Before the experiments, agarose was melted in the microwave and kept in a water bath at 65°C. To make the bottom layer, agarose solution was mixed with 10% serum-containing medium to the final concentration of 0.7% agarose. 1.5ml of this was laid to make the bottom layer. While this layer was solidifying, cells were trypsinised and made into a single-cell suspension. For the top layer, for each well, 3×10^4^ cells were mixed with the agarose and serum-containing media to the final agarose concentration of 0.35%. 1.5ml of this cell suspension was quickly dispensed on top. Finally, 2.5ml of 10% serum-containing media were pipetted on top. The cells were grown for 10 days with the media were changed every 2 days. Five random fields of cell spheroids per condition were imaged with a standard brightfield microscope. Image quantification was done manually by outlining the spheroids with the Freehand selection tool from ImageJ and the area of the spheroid were calculated. Only in-focused spheroids were measured.

### Cloning and primers

Making the internal GFP or mCherry CYRI-A construct: Mouse CYRI-A cDNA was cloned into the pcDNA3.1(+) vector (a kind gift from Dr Douglas Strathdee) using standard cloning technique. HindIII and BamHI restriction sites were inserted just after codon coding for Proline 16 using the Q5-insertion mutagenesis (NEB, #E0554) according to the protocol from the manufacturer. GFP or mCherry were PCR out from pEGFP-C1 or pmCherry-N1 vector before being inserted internally into CYRI-A.

All primers were ordered from ThermoFisher Scientifics:

PCR CYRI-A

5′-TCAACGGCTAGCATGGGAAACTTGCTCAAAGTC-3′

5′-TCAACGCTCGAGTTACTGAAGCATCGCTCGAATCTG-3′

Insert restriction sites

5′-AAGCTTGCGGGATCCCACTTCTTCCTGGATTTTGAAAATG-3′

5′-GGATCCCGCAAGCTTTGGATAATTCTCAATTTCCC-3′

PCR GFP or mCherry

5′-TCAACGAAGCTTGTGAGCAAGGGCGAGGAGCTG-3′

5′-TCAACGGGATCCCTTGTACAGCTCGTCCATGCC-3′

5′-TCAACGAAGCTTGTGAGCAAGGGCGAGGAGG-3′

Insert Kozak sequence

5′-GCCACCATGGGAAACTTGCTCAAAGTCC-3′

5′-GCTAGCCAGCTTGGGTCT-3′

The same strategy is used to clone for P17-GFP-CYRI-B but the restriction sites are XHoI and NheI just after the codon code for Proline 17.

5′-CTCGAGGCGGCTAGCAATTTTTTCCTTGATTTTGA-3′

5′-GCTAGCCGCCTCGAGTGGCCCCTGCTCAAGGTCTG-3′

Primers for GFP-CYRI-A (1-319) and GFP-CYRI-A (29-319)

5′-TCAACGCTCGAGTGATGGGAAACTTGCTCAAAGTC-3′

5′-CATGCAGGATCCTTACTGAAGCATCGCTCGAATCTG-3′

5′-TCAACGCTCGAGTGGAAGGAGAGCGAGAGATATGG-3′

5′-CATGCAGGATCCTTATCGAATCTGTTTGGAAGTCGA-3′

Primers for the mVenus-bi-cistronic vector of CYRI-A

5′-TCAACGGCTAGCGCCACCATGGGAAACTTGCTCAAAGTC-3′

5′-TCAACGGGATCCTTACTGAAGCATCGCTCGAATCTG-3′

Primers for CYRI-A-FLAG

5′-TCAACGGGATCCGCCACCATGGGAAACTTGCTCAAAGTC-3′

5′-

TCAACGCTCGAGTTACTTGTCGTCATCGTCTTTGTAGTCCTGAAGCATCGCTCGAATCTG-3′

Primers for MBP-CYRI-A

5′-CGTATTGGATCCATG GGAAACTTG CTCAAAGTC-3′

5′-CGTATTGAAGCTTCTACTGAAGCATCGCTCGAAT-3

GFP-PH-Grp1 was a kind gift from Dr David Bryant.

### Addgene plasmids

mApple-Alpha-5-Integrin-12 was a gift from Michael Davidson (Addgene plasmid # 54864; http://n2t.net/addgene:54864; RRID:Addgene_54864) mCherry-Clathrin LC-15 was a gift from Michael Davidson (Addgene plasmid # 55019; http://n2t.net/addgene:55019; RRID:Addgene_55019) (Rizzo et al., 2009) pcDNA3/hArf1(WT)-mCherry was a gift from Kazuhisa Nakayama (Addgene plasmid # 79419; http://n2t.net/addgene:79419; RRID:Addgene_79419) (Makyio et al., 2012) CAV1-mCherry was a gift from Ari Helenius (Addgene plasmid # 27705; http://n2t.net/addgene:27705; RRID:Addgene_27705) (Hayer et al., 2010) mCherry-Rab5a-7 was a gift from Michael Davidson (Addgene plasmid # 55126; http://n2t.net/addgene:55126; RRID:Addgene_55126) PH-Btk-GFP was a gift from Tamas Balla (Addgene plasmid # 51463; http://n2t.net/addgene:51463; RRID:Addgene_51463) (Varnai and Balla, 1998)

### Antibodies

Anti CYRI-A antibody (Sigma-Aldrich, not in production, western blotting), Anti CYRI-B antibody (Sigma-Aldrich, #HPA009076, western blotting), Anti GAPDH (Millipore, #MAB374, western blotting), Anti Tubulin DM1A (Sigma-Aldrich, #T6199, western blotting), Anti GST (CST, #2622, western blotting), Anti GFP (CST, #2955, western blotting), Phalloidin 488 (Molecular Probe, #A12379), Anti-Integrin beta 1 antibody [12G10] (Abcam, #ab30394, immunofluorescent and western blotting), Recombinant Anti-Integrin alpha 5 antibody [EPR7854] (Abcam, #ab150361, immunofluorescent and western blotting), Anti-Integrin alpha5 (Preservative Free) Antibody, clone SNAKA51 (Millipore, #MABT201, immunofluorescent, internalisation assay and flow cytometry), Alexa Fluor 488 anti-human CD29 Antibody TS2/16 (BioLegend, #303015, flow cytometry), Alexa Fluor 488 anti-mouse CD49e Antibody (BioLegend, #103810, flow cytometry), Anti-MMP-14 Antibody, catalytic domain, clone LEM-2/63.1 (Millipore, #MAB3329, flow cytometry).

### Statistical analysis

All statistical analyses are performed with Prism 7. All data points are pooled from independent experiments or cells and then performed the appropriate statistical test. Normality is assessed by inspecting the shape of the distribution. Parametric tests are used because they are less affected by deviation from normality distribution and are more robust than non-parametric tests (McDonald, J.H. 2014. Handbook of Biological Statistics (3rd ed.). Sparky House Publishing, Baltimore, Maryland).

## Acknowledgements

We thank Dr Sue Burchill and Andrea Berry from the Leeds Institute of Cancer and Pathology, Cancer Research UK Leeds Centre for providing us with the A-673 cell line, Dr David Bryant and Dr Alvaro Roman Fernandez from the Cancer Research UK Beatson Institute and the University of Glasgow for sharing with us the PH-Grp1 construct. We thank Cancer Research UK for core funding (A17196 and A31287) to LMM (A24452) and SI (A19257). We thank Margaret O’Prey, David Strachan and Tom Gilbey for their help with confocal microscopy, Incucyte and flow cytometry, respectively. We thank Jillian Murray for the help with making CDM plates. We thank the Beatson central service, the molecular service and histology service for their help with DNA sequencing and histology slide processing and analysis, respectively.

---

**Figure S1.**
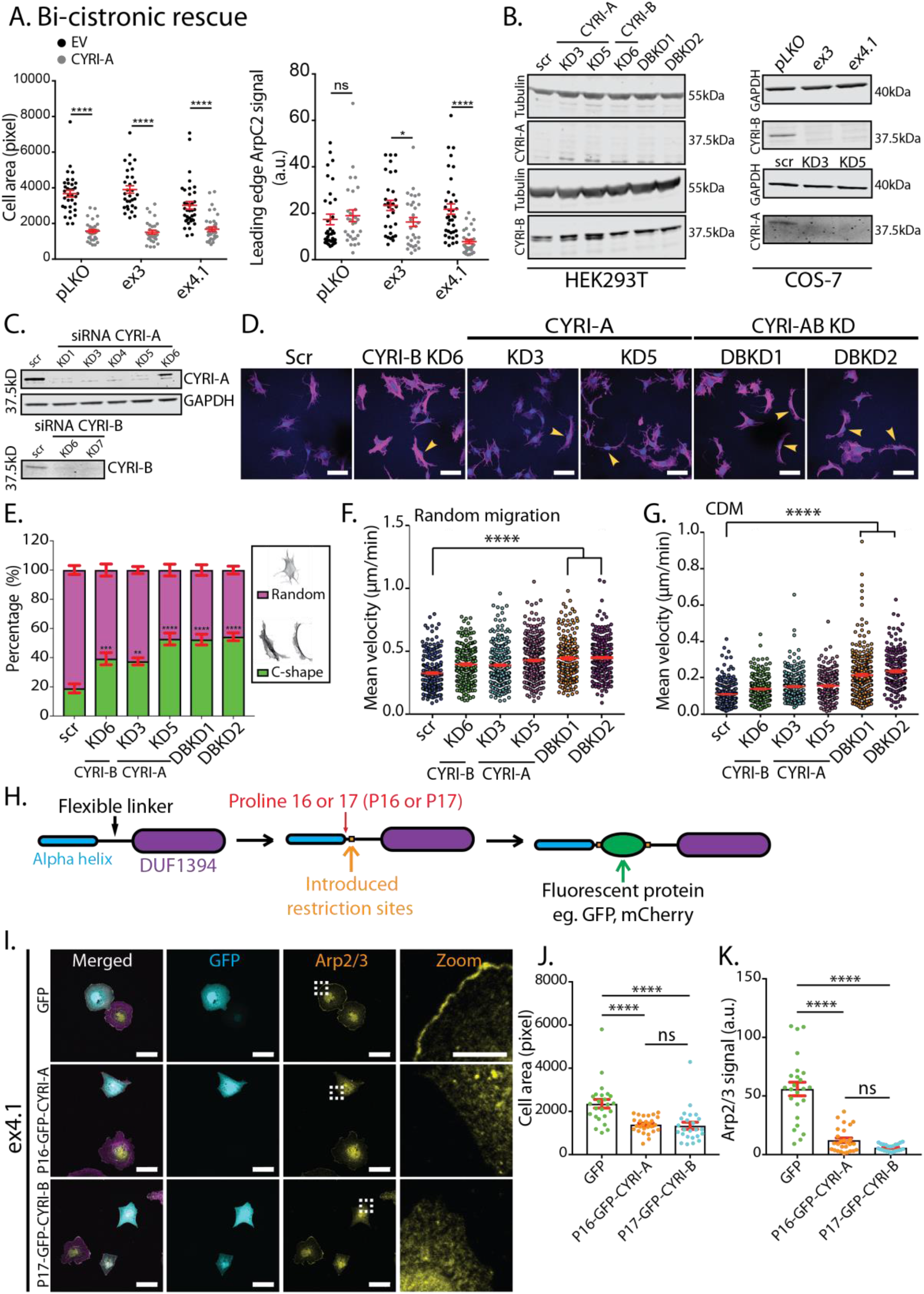
CYRI-A cooperates with CYRI-B to restrict lamellipodia. For A-G, pLKO = control CRISPR line, ex3 and ex4.1 = CYRI-B CRISPR knockout cell lines, scr = siRNA scramble, KD3 and KD5 = CYRI-A siRNA knockdowns, KD6 = CYRI-B siRNA knockdown, DBKD1/2 = double CYRI-A/B knockdown (KD3+KD6 or KD5+KD6, respectively). A. Quantification of cell area (left) and the average Arp2/3 signal intensity around the cell perimeter in COS-7 CRISPR cells (ex3 and ex4.1) (black dots) and rescued with untagged CYRI-A (grey dots). Data from 3 independent experiments, statistical analysis using unpaired t-test. Mean ± SEM. B. Western blot of control or knockdown HEK293T cells (left panels). Western blot (right panels) of endogenous CYRI-A and CYRI-B in COS-7 cells and CRISPR KO. Tubulin and GAPDH are loading control. C. Western blot of endogenous CYRI-A and CYRI-B in control or knockdown A-673 cells. GAPDH is loading control. D. Representative immunofluorescence images of control scrambled (Scr), single CYRI-A or CYRI-B and CYRI-A/B double knockdown (DBKD) A-673 stained for F-actin (magenta) and nuclei (blue). Yellow arrowheads point to C-shape cells. Scale bar = 50μm. E. Quantification of the number of C-shaped cells in D. Data from 3 independent experiments with at least 50 cells per experiment. ANOVA with multiple comparisons. Mean ± SEM. **p<0.01, ***p<0.001, ****p<0.0001. F. G. Quantification of the mean velocity of control, single and CYRI-A/B double knockdown A-673 cells plated for 2D fibronectin (F) or 3D cell-derived matrix (CDM) (G). Data from 3 independent experiments. ANOVA with Tukey’s multiple comparison test. Mean ± SEM. ****p<0.0001. H. Schematic representation of the cloning strategy used to create internally tagged P16-GFP-CYRI-A and P17-GFP-CYRI-B. I. Immunofluorescence images of control or CYRI-B CRISPR knockout (ex4.1) COS-7 transfected with either P16-GFP-CYRI-A or P17-GFP-CYRI-B (cyan) and stained for Arp2/3 (yellow) and F-actin (magenta). Scale bar = 20μm for full-size image or 10μm for zoom. J-K. Quantification of the cell area and the average Arp2/3 signal at the cell periphery of CYRI-B CRISPR COS-7 cells ex4.1 expressing control GFP, P16-GFP-CYRI-A or P17-GFP-CYRI-B. Data from 3 independent experiments with Tukey’s multiple comparison test with at least 10 cells per experiment. ns = p>0.05, ****p<0.0001.

**Figure S2.**
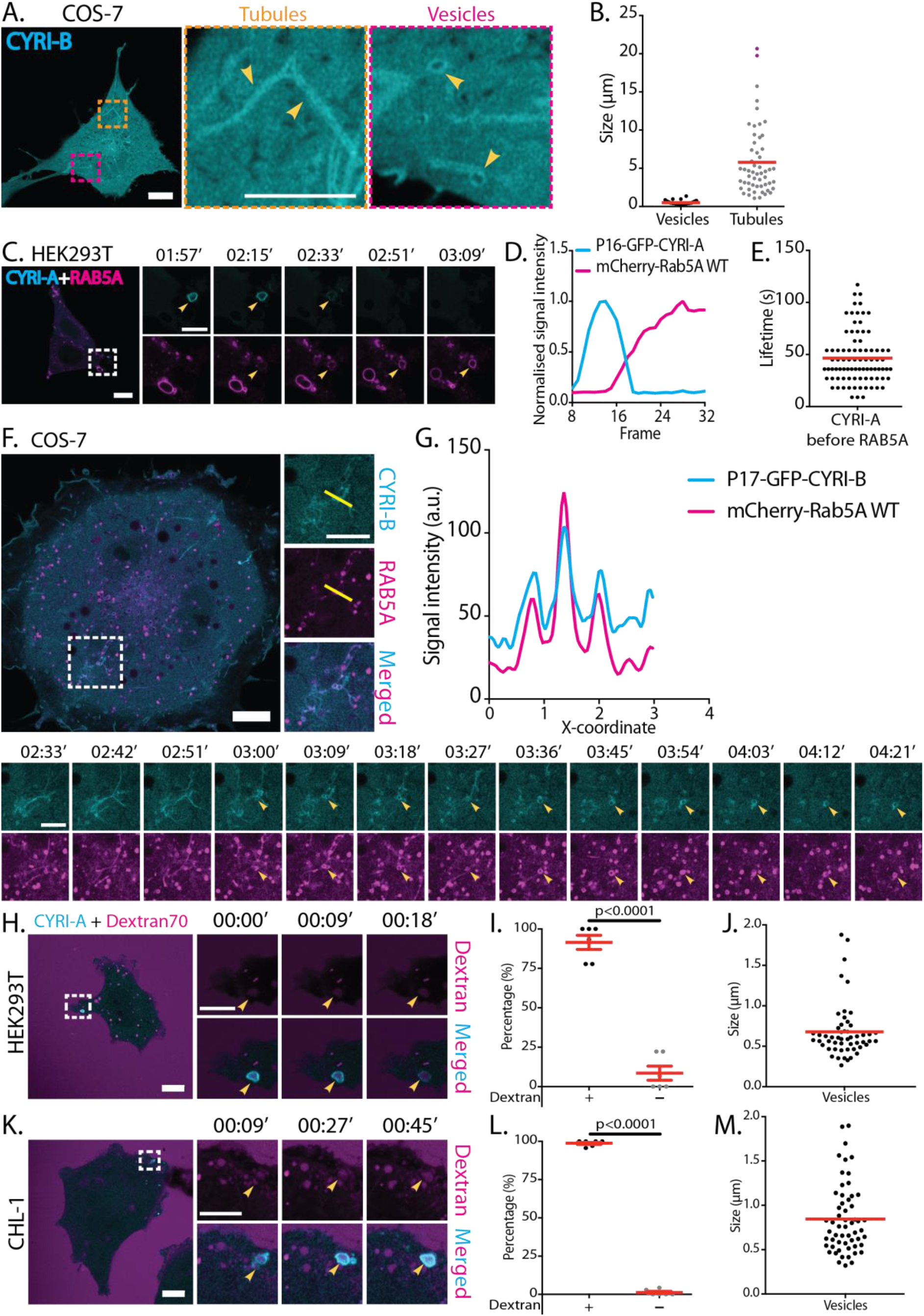
Dynamic localisation of CYRI proteins on macropinosomes and tubulovesicular structures. A-B. Representative images of live COS-7 cells expressing the P17-GFP-CYRI-B (scale bar = 10μm). Tubular and vesicular structures are highlighted in zoomed panels and quantification of their sizes shown in B. (vesicles N=16 events in 3 cells, tubules N=22 events in 4 cells) (scale bar = 5μm). See Video 21. C-E. Time sequence of live HEK293T cells (scale bar = 10μm) co-expressing P16-GFP-CYRI-A (cyan) and mCherry-RAB5A WT (magenta). Arrowhead points to vesicular structures (C) (scale bar = 5μm). Dynamics of each protein is reported by its normalised intensity plot (D) and lifetime (N=84 events in 7 cells) (E). See Video 22. F-G. COS-7 cells (scale bar = 10μm) co-expressing P16-GFP-CYRI-B (cyan) and mCherry-RAB5A WT (magenta). Time sequence corresponding to the white dotted square area is shown in the bottom panel (scale bar = 5μm). Arrowhead points to tubular and vesicular structures and intensity profile along the yellow line is plotted in G. See Video 23. H-M. Time sequence images HEK293T (H) and CHL-1 cells (K) expressing P16-GFP-CYRI-A and incubated with dextran 70kDa (scale bar = 10μm). Yellow arrowheads indicate macropinocytic events positive for both CYRI-A and dextran signals (scale bar = 5μm). Quantification showing the majority of CYRI-A positive vesicles are also dextran-positive in HEK293T (I) (88%, N=6 cells) and CHL-1 (L) (100%, N=6 cells) and their sizes (J and M) (N=53 events in 6 cells in HEK293T, N=57 events in 6 cells in CHL-1). See Video 24-25. Red line indicates the average value.

**Figure S3.**
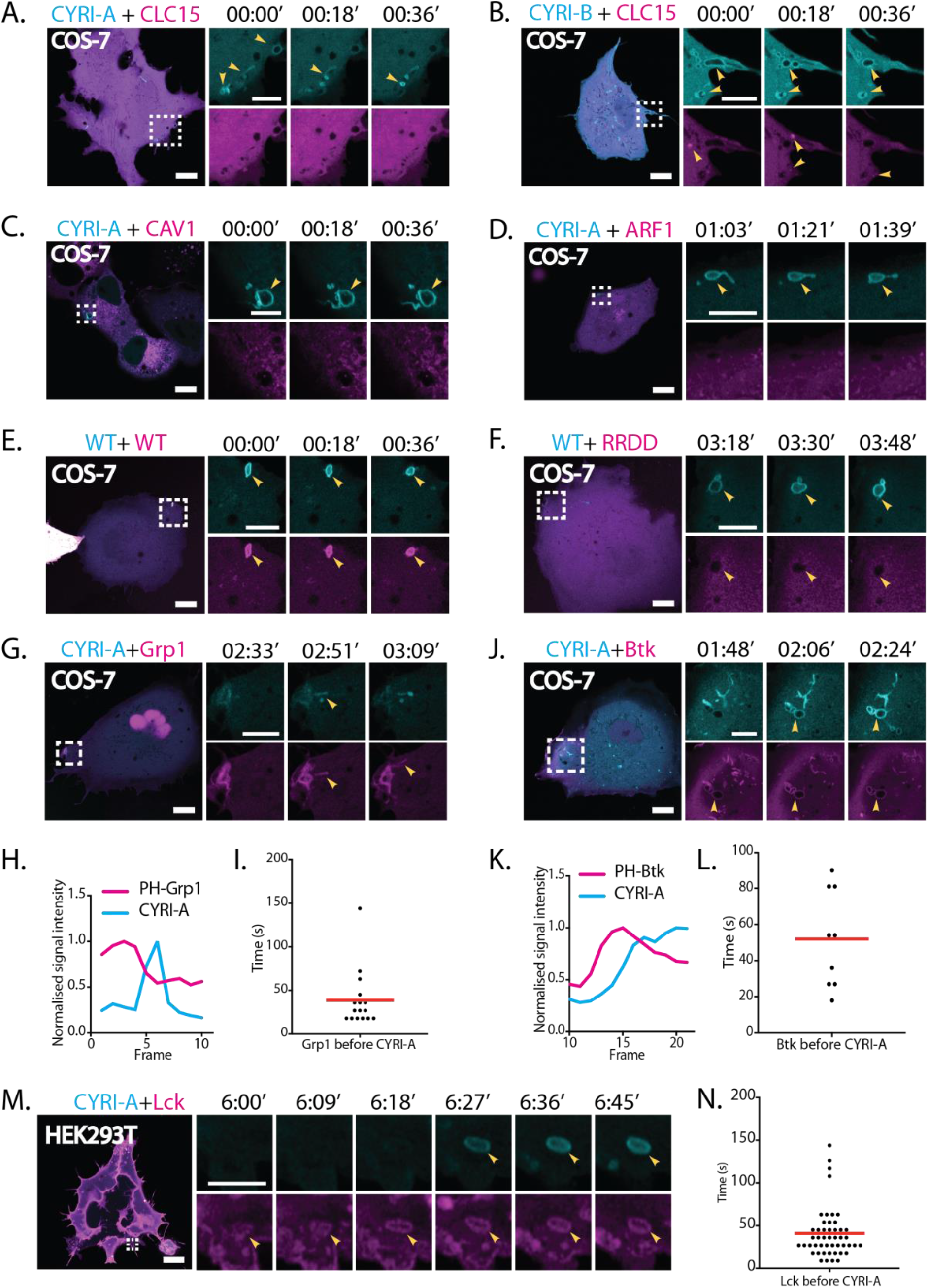
CYRI-A does not co-localise with endocytic markers, but co-localises with markers of PIP3. A-D. Time sequence images of COS-7 cells expressing P16-GFP-CYRI-A or P17-GFP-CYRI-B (cyan) and either mCherry-tagged CLC15 (for Clathrin Light Chain 15, magenta) (A), Caveolin-1(C, magenta) or ARF1(D, magenta). Scale bar = 10μm for full-sized images and 5μm for the zooms. See Videos 26-29. E-F. Time sequence images of live COS-7 cells expressing mCherry-WT or RRDD mutant P16-GFP-CYRI-A (magenta) and GFP-WT P16-GFP-CYRI-A (cyan). See Videos 30-31. G-L. COS-7 co-expressing P16-GFP-CYRI-A (cyan) and two independent PIP3 reporters (magenta) PH-Grp1 (G-I) or PH-Btk (J-L) (N=31 events in 3 cells for Grp1 and N=9 events in 1 cell for Btk). Red line represents the average value. Scale bar = 10μm for full-size image and 5μm for the zooms. See Videos 32-33. M-N. Time sequence images of HEK293T expressing P16-GFP-CYRI-A and mScarlet-Lck (labelling the plasma membrane). The time Lck resides on the vesicles before CYRI-A is recruited is quantified in N (N=48 events in 10 cells). Scale bar = 10μm for full-size image and 3μm for the zooms. See Video 34.

**Figure S4.**
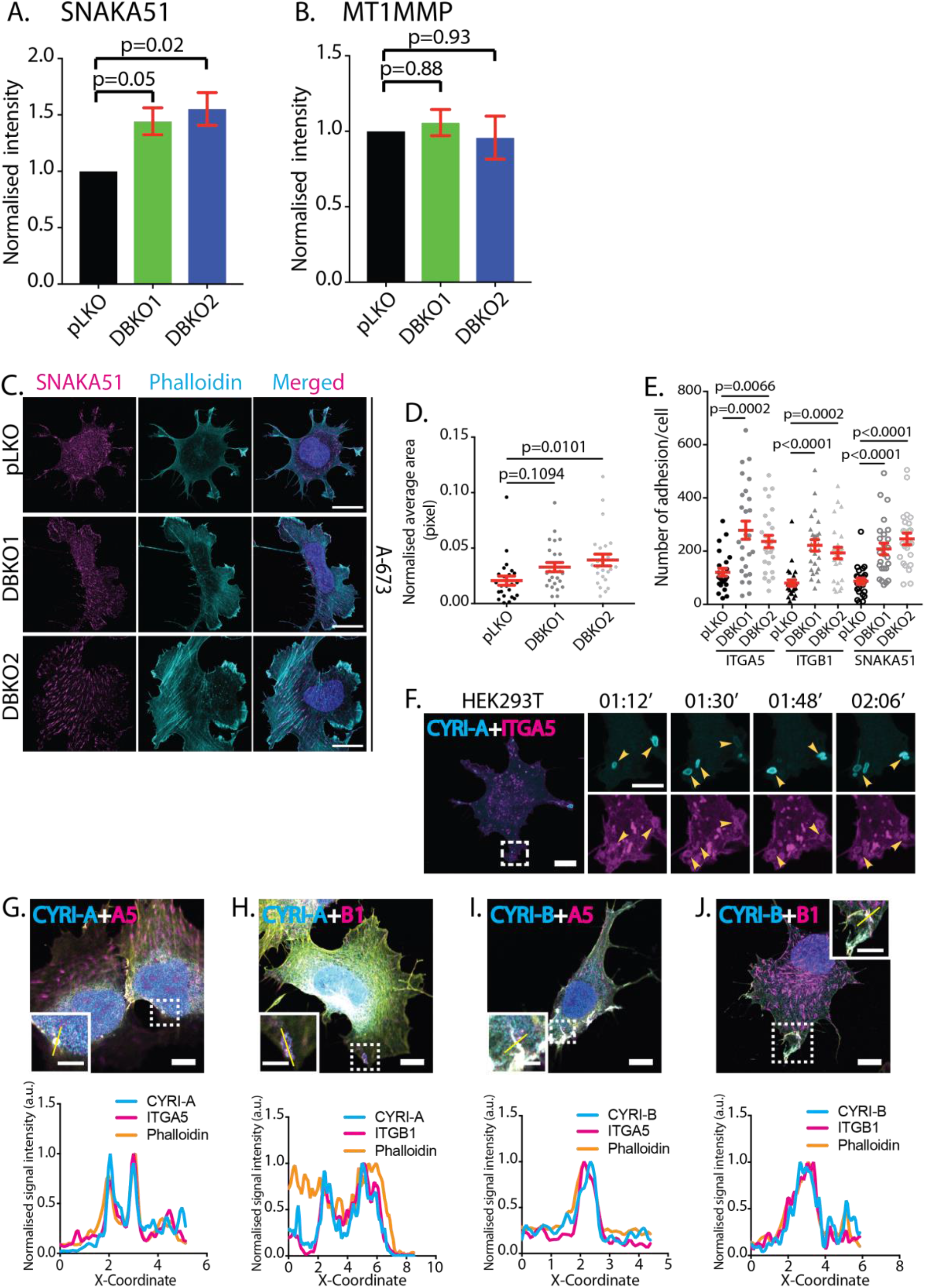
Integrin α5β1 is a major target of micropinocytosis orchestrated by CYRI proteins. A-B. Flow cytometry analysis of surface expression of active integrin α5, detected using the SNAKA51 antibody (A) and MT1-MMP (B) comparing between the control pLKO and CYRI-A/B double knockout DBKO A673 cells. Data from 3 independent experiments. Statistical analysis using ANOVA with Tukey’s multiple comparison test. C-E. Immunofluorescence images of the control pLKO and DBKO A-673 cells stained for active integrin α5 (magenta) and actin (cyan). The average area of integrin clusters or the number of clusters per cell are quantified in D and E. Data from 3 independent experiments with at least 10 cells per experiment. Mean ± SEM. Statistical analysis use ANOVA with Tukey’s multiple comparison test. Scale bars = 20μm. F. Time sequence images of HEK293T expressing P16-GFP-CYRI-A (cyan) and mApple-integrin α5 (magenta) showing integrin α5 signal present on CYRI-A-positive vesicles. Scale bar = 10μm for full-sized image and 5μm for zooms. See Video 35. G-J. Immunofluorescence images of endogenous integrins α5 and β1 in A673 cells with P16-GFP-CYRI-A or P17-GFP-CYRI-B along with actin (yellow). Graphs show the colocalization of CYRI-A, integrins and filamentous actin (phalloidin) on the vesicles. Scale bars = 10μm. In C,G-J: DAPI for DNA.

**Figure S5.**
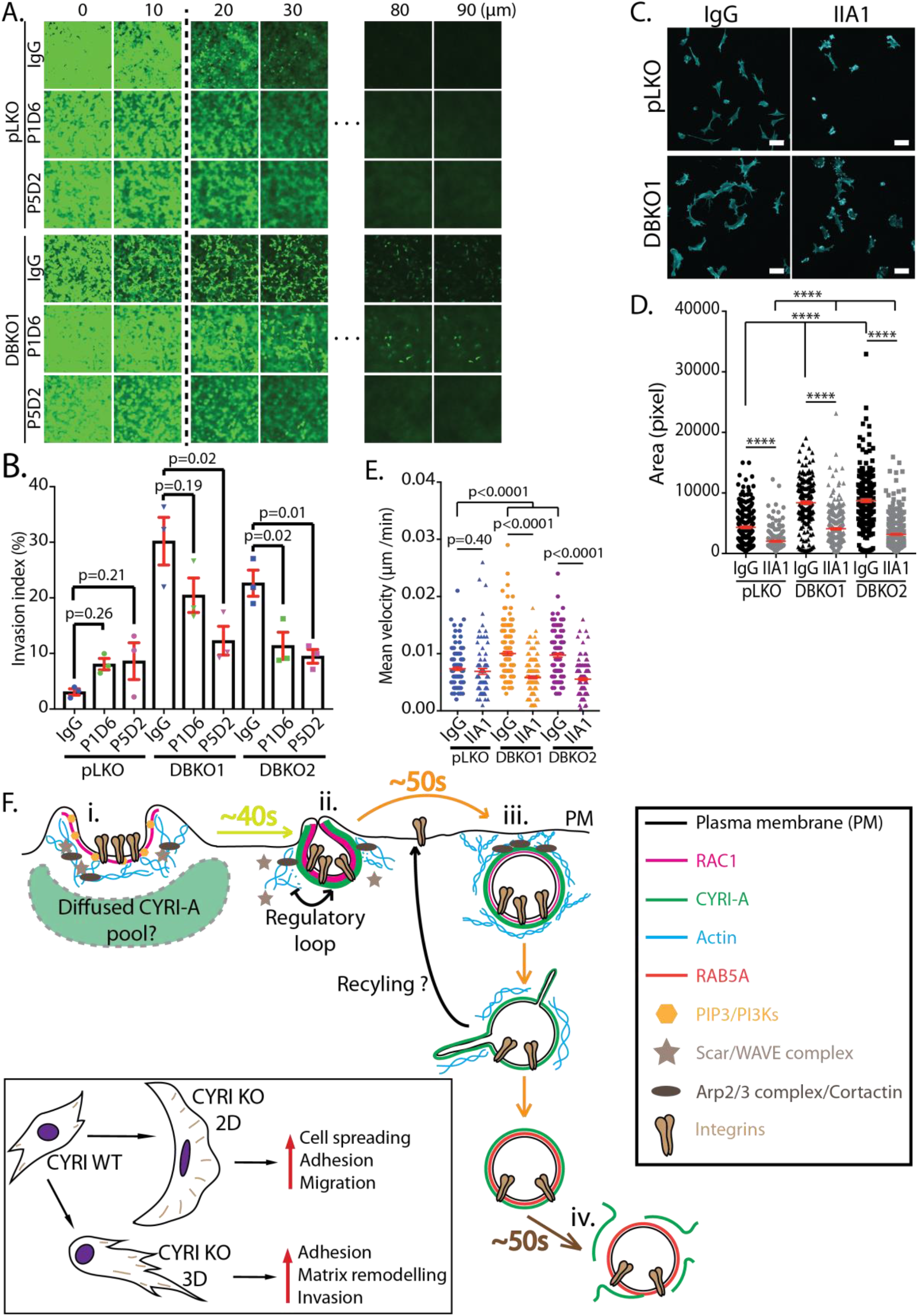
Failure of integrin α5β1 macropinocytosis enhances cell invasion and working model of CYRI function. A-B. Inverted invasion assay of control pLKO and CYRI DBKO A-673 cells with collagen-Matrigel-fibronectin (CMF) plug treated with IgG (negative control), P1D6 (α5-blocking antibody) or P5D2 (β1-blocking antibody) (A). The depth of invasion beyond 10 microns was quantified as an invasion index shown in B. Data from 3 independent experiments. Mean ± SEM. Statistical analysis using ANOVA with Tukey’s multiple comparison test. C-D. Immunofluorescence images of control pLKO or CYRI-A/B double knockout DBKO A-673 cells treated with 5μg/ml of control IgG antibody or α5β1-complex-blocking antibody IIA1 and stained for F-actin. Cell area was measured and plotted in D. Data from 3 independent experiments. Mean ± SEM. Statistical analysis using unpaired t-test or ANOVA with Tukey’s multiple comparison test where appropriate. Scale bar = 50μm. E. Treating DBKO A-673 with integrin blocking antibody IIA1 shows the reduction in the migration ability on 2D. Data from 3 independent experiments. Statistical analysis using unpaired t-test or ANOVA with Tukey’s multiple comparison test where appropriate. F. The current working model: i. RAC1, the Scar/WAVE complex, the Arp2/3 complex and actin drives the formation of macropinocytic cup. Ii. RAC1 activity increases leads to the recruitment of CYRI-A from a diffuse pool. Iii. CYRI-A is recruited to the nascent macropinosome by active RAC1, where it dampens down RAC1 activity, terminates actin polymerisation and allows for the completion of macropinosomes, taking surface integrins into the cytoplasm. iv. CYRI-A slowly disappears while RAB5A starts to appear. Cancer cells lacking both CYRIs retain more integrins on their surface, which can help them adhere to their surrounding matrix to assist their migration and invasion.

## Supplementary videos

Video 1 (Related to Fig. 4 A).

Airy-scan video of HEK293T cells expressing P16-GFP-CYRI-A (cyan). Arrows indicate vesicles. Acquisition at 9s/frame and play back at 10 fps.

Video 2 (Related to Fig. 4 D).

Airy-scan video of HEK293T cells expressing P16-GFP-CYRI-A (cyan). Arrows indicate vesicles and the diffuse CYRI-A pool. Acquisition at 9s/frame and play back at 10 fps.

Video 3 (Related to Fig. 4 E).

Airy-scan video of COS-7 cells expressing P16-GFP-CYRI-A (cyan) and mCherry-RAB5A WT (magenta). Arrows indicate the vesicle. Acquisition at 9s/frame and play back at 10 fps.

Video 4 (Related to Fig. 4 H).

Airy-scan video of COS-7 cells expressing P16-GFP-CYRI-A (cyan) and dextran 70kDa (magenta). Arrows indicate the vesicle containing the dextran. Acquisition at 9s/frame and play back at 10 fps.

Video 5 (Related to Fig. 5 A).

Airy-scan video of COS-7 cells expressing GFP (cyan) and LifeAct-RFP (magenta). Acquisition at 9s/frame and play back at 10 fps.

Video 6 (Related to Fig. 5 A).

Airy-scan video of COS-7 cells expressing P16-GFP-CYRI-A (cyan) and LifeAct-RFP (magenta). Arrows indicate the vesicles. Acquisition at 9s/frame and play back at 10 fps.

Video 7 (Related to Fig. 5 B).

Airy-scan video of a zoom of COS-7 cells expressing P16-GFP-CYRI-A (cyan) and LifeAct-RFP (magenta). Arrows indicate the vesicles with actin around. Acquisition at 9s/frame and play back at 10 fps.

Video 8 (Related to Fig. 5 C).

Airy-scan video of a zoom of HEK293T cells expressing P16-GFP-CYRI-A (cyan) and LifeAct-RFP (magenta). Arrows indicate the vesicle with actin around. Acquisition at 9s/frame and play back at 10 fps.

Video 9 (Related to Fig. 5 E).

Airy-scan video of DBKD COS-7 cells expressing GFP (cyan) and LifeAct-RFP (magenta). Acquisition at 9s/frame and play back at 10 fps.

Video 10 (Related to Fig. 5 E).

Airy-scan video of DBKD COS-7 cells expressing P16-GFP-CYRI-A (cyan) and LifeAct-RFP (magenta). Arrows indicate the vesicles. Acquisition at 9s/frame and play back at 10 fps.

Video 11 (Related to Fig. 6 A).

Airy-scan video of HEK293T cells expressing P16-mCherry-CYRI-A (cyan) and GFP-RAC1 WT (magenta). Arrows indicate the vesicles. Acquisition at 9s/frame and play back at 10 fps.

Video 12 (Related to Fig. 6 D).

Airy-scan video of HEK293T cells expressing P16-mCherry-CYRI-A (cyan) and CFP-PBD (magenta). Arrows indicate the vesicle. Acquisition at 9s/frame and play back at 10 fps.

Video 13 (Related to Fig. 6 F).

Airy-scan video of HEK293T cells expressing P16-GFP-CYRI-A WT (cyan) and P16-mCherry-CYRI-A WT (magenta). Arrows indicate the vesicle. Acquisition at 9s/frame and play back at 10 fps.

Video 14 (Related to Fig. 6 I).

Airy-scan video of HEK293T cells expressing P16-GFP-CYRI-A WT (cyan) and P16-mCherry-CYRI-A RRDD (magenta). Arrows indicate the vesicle. Acquisition at 9s/frame and play back at 10 fps.

Video 15 (Related to Fig. 7 A).

Airy-scan video of HEK293T cells expressing P16-mCherry-CYRI-A (cyan) and GFP-GRP1 (magenta). Arrows indicate the vesicle. Acquisition at 9s/frame and play back at 10 fps.

Video 16 (Related to Fig. 7 D).

Airy-scan video of HEK293T cells expressing P16-mCherry-CYRI-A (cyan) and GFP-Btk (magenta). Arrows indicate the vesicle. Acquisition at 9s/frame and play back at 10 fps.

Video 17 (Related to Fig. 7 G).

Example 1 of an airy-scan video of COS-7 cells expressing P16-mCherry-CYRI-A (cyan) and treated with 20μM LY294002 (as indicated). Arrows indicate the vesicles. Acquisition at 30s/frame and play back at 7 fps.

Video 18 (Related to Fig. 7 G).

Example 2 of an airy-scan video of COS-7 cells expressing P16-mCherry-CYRI-A (cyan) and treated with 20μM LY294002 (as indicated). Arrows indicate the vesicles. Acquisition at 30s/frame and play back at 7 fps.

Video 19 (Related to Fig. 9 A).

Airy-scan video of COS-7 cells expressing P16-GFP-CYRI-A (cyan) and mApple-ITGA5 (magenta). Arrows indicate the vesicles. Acquisition at 9s/frame and play back at 10 fps.

Video 20 (Related to Fig. 9 B).

Airy-scan video of COS-7 cells expressing P17-GFP-CYRI-B (cyan) and mApple-ITGA5 (magenta). Arrows indicate the vesicles. Acquisition at 9s/frame and play back at 10 fps.

Video 21 (Related to SFig. 2 A).

Airy-scan video of COS-7 cells expressing P17-GFP-CYRI-B (cyan). Acquisition at 9s/frame and play back at 10 fps.

Video 22 (Related to SFig. 2 C).

Airy-scan video of HEK293T cells expressing P16-GFP-CYRI-A (cyan) and mCherry-RAB5A WT (magenta). Arrows indicate the vesicles. Acquisition at 9s/frame and play back at 10 fps.

Video 23 (Related to SFig. 2 F).

Airy-scan video of COS-7 cells expressing P17-GFP-CYRI-B (cyan) and mCherry-RAB5A WT (magenta). Acquisition at 9s/frame and play back at 10 fps.

Video 24 (Related to SFig. 2 H).

Airy-scan video of CHL-1 cells expressing P16-GFP-CYRI-A (cyan) and dextran 70kDa (magenta). Acquisition at 9s/frame and play back at 10 fps.

Video 25 (Related to SFig. 2 H).

Airy-scan video of HEK293T cells expressing P16-GFP-CYRI-A (cyan) and dextran 70kDa (magenta). Acquisition at 9s/frame and play back at 10 fps.

Video 26 (Related to SFig. 3 A).

Airy-scan video of COS-7 cells expressing P16-GFP-CYRI-A (cyan) and mCherry-CLC15 (clathrin) (magenta). Acquisition at 9s/frame and play back at 10 fps.

Video 27 (Related to SFig. 3 B).

Airy-scan video of COS-7 cells expressing P17-GFP-CYRI-A (cyan) and mCherry-CLC15 (clathrin) (magenta). Acquisition at 9s/frame and play back at 10 fps.

Video 28 (Related to SFig. 3 C).

Airy-scan video of COS-7 cells expressing P16-GFP-CYRI-A (cyan) and mCherry-Cav1 (caveolin) (magenta). Acquisition at 9s/frame and play back at 10 fps.

Video 29 (Related to SFig. 3 D).

Airy-scan video of COS-7 cells expressing P16-GFP-CYRI-A (cyan) and mCherry-ARF1 (magenta). Acquisition at 9s/frame and play back at 10 fps.

Video 30 (Related to SFig. 3 E).

Airy-scan video of COS-7 cells expressing P16-GFP-CYRI-A WT (cyan) and P16-mCherry-CYRI-A WT (magenta). Acquisition at 9s/frame and play back at 10 fps.

Video 31 (Related to SFig. 3 F).

Airy-scan video of COS-7 cells expressing P16-GFP-CYRI-A WT (cyan) and P16-mCherry-CYRI-A RRDD (magenta). Acquisition at 9s/frame and play back at 10 fps.

Video 32 (Related to SFig. 3 G).

Airy-scan video of COS-7 cells expressing P16-mCherry-CYRI-A (cyan) and GFP-GRP1 (magenta). Arrows indicate the vesicle. Acquisition at 9s/frame and play back at 10 fps.

Video 33 (Related to SFig. 3 J).

Airy-scan video of COS-7 cells expressing PI6-mCherry-CYRI-A (cyan) and GFP-Btk1 (magenta). Arrows indicate the vesicle. Acquisition at 9s/frame and play back at 10 fps.

Video 34 (Related to SFig. 3 J).

Airy-scan video of HEK293T cells expressing P16-GFP-CYRI-A (cyan) and mScarlet-Lck (magenta). Arrows indicate the vesicle. Acquisition at 9s/frame and play back at 10 fps.

Video 35 (Related to SFig. 4 F).

Airy-scan video of HEK293T cells expressing P16-GFP-CYRI-A (cyan) and mApple-ITGA5 (magenta). Arrows indicate the vesicles. Acquisition at 9s/frame and play back at 10 fps.

## References

Alanko, J., A. Mai, G. Jacquemet, K. Schauer, R. Kaukonen, M. Saari, B. Goud, and J. Ivaska. 2015. Integrin endosomal signalling suppresses anoikis. Nat Cell Biol. 17:1412–1421.

Araki, N., Y. Egami, Y. Watanabe, and T. Hatae. 2007. Phosphoinositide metabolism during membrane ruffling and macropinosome formation in EGF-stimulated A431 cells. Exp Cell Res. 313:1496–1507.

Araki, N., M.T. Johnson, and J.A. Swanson. 1996. A role for phosphoinositide 3-kinase in the completion of macropinocytosis and phagocytosis by macrophages. J Cell Biol. 135:1249–1260.

Azevedo, C.M.S., R.A. Machado, H. Martelli-Junior, S.R.A. Reis, D.C. Persuhn, R.D. Coletta, and A. Rangel. 2020. Exploring GRHL3 polymorphisms and SNP-SNP interactions in the risk of non-syndromic oral clefts in the Brazilian population. Oral Dis. 26:145–151.

Bianchi-Smiraglia, A., S. Paesante, and A.V. Bakin. 2013. Integrin beta5 contributes to the tumorigenic potential of breast cancer cells through the Src-FAK and MEK-ERK signaling pathways. Oncogene. 32:3049–3058.

Bloomfield, G., and R.R. Kay. 2016. Uses and abuses of macropinocytosis. J Cell Sci. 129:2697–2705.

Bohdanowicz, M., and S. Grinstein. 2013. Role of phospholipids in endocytosis, phagocytosis, and macropinocytosis. Physiol Rev. 93:69–106.

Buckley, C.M., and J.S. King. 2017. Drinking problems: mechanisms of macropinosome formation and maturation. FEBS J. 284:3778–3790.

Campa, C.C., E. Ciraolo, A. Ghigo, G. Germena, and E. Hirsch. 2015. Crossroads of Pl3K and Rac pathways. Small GTPases. 6:71–80.

Cannon, A.C., C. Uribe-Alvarez, and J. Chernoff. 2020. RAC1 as a Therapeutic Target in Malignant Melanoma. Trends Cancer. 6:478–488.

Canton, J. 2018. Macropinocytosis: New Insights Into Its Underappreciated Role in Innate Immune Cell Surveillance. Front Immunol. 9:2286.

Carlsson, A.E. 2018. Membrane bending by actin polymerization. Curr Opin Cell Biol. 50:1–7.

Caswell, P.T., M. Chan, A.J. Lindsay, M.W. McCaffrey, D. Boettiger, and J.C. Norman. 2008. Rab-coupling protein coordinates recycling of alpha5beta1 integrin and EGFR1 to promote cell migration in 3D microenvironments. J Cell Biol. 183:143–155.

Caswell, P.T., S. Vadrevu, and J.C. Norman. 2009. Integrins: masters and slaves of endocytic transport. Nat Rev Mol Cell Biol. 10:843–853.

Chattaragada, M.S., C. Riganti, M. Sassoe, M. Principe, M.M. Santamorena, C. Roux, C. Curcio, A. Evangelista, P. Allavena, R. Salvia, B. Rusev, A. Scarpa, P. Cappello, and F. Novelli. 2018. FAM49B, a novel regulator of mitochondrial function and integrity that suppresses tumor metastasis. Oncogene. 37:697–709.

Chen, C., Q. Guo, J. Shi, X. Jiao, K. Lv, X. Liu, Y. Jiang, X. Hui, and T. Song. 2018. Genetic variants of MGMT, RHPN2, and FAM49A contributed to susceptibility of nonsyndromic orofacial clefts in a Chinese population. J Oral Pathol Med. 47:796–801.

Chertkova, A.O., M. Mastop, M. Postma, N. van Bommel, S. van der Niet, K.L. Batenburg, L. Joosen, T.W.J. Gadella, Y. Okada, and J. Goedhart. 2020. Robust and Bright Genetically Encoded Fluorescent Markers for Highlighting Structures and Compartments in Mammalian Cells. BioRXIV.

Clark, K., R. Pankov, M.A. Travis, J.A. Askari, A.P. Mould, S.E. Craig, P. Newham, K. M. Yamada, and M.J. Humphries. 2005. A specific alpha5beta1-integrin conformation promotes directional integrin translocation and fibronectin matrix formation. J Cell Sci. 118:291–300.

Commisso, C., R.J. Flinn, and D. Bar-Sagi. 2014. Determining the macropinocytic index of cells through a quantitative image-based assay. Nat Protoc. 9:182192.

Condon, N.D., J.M. Heddleston, T.L. Chew, L. Luo, P.S. McPherson, M.S. Ioannou, L. Hodgson, J.L. Stow, and A.A. Wall. 2018. Macropinosome formation by tent pole ruffling in macrophages. J Cell Biol. 217:3873–3885.

Cooper, J., and F.G. Giancotti. 2019. Integrin Signaling in Cancer: Mechanotransduction, Stemness, Epithelial Plasticity, and Therapeutic Resistance. Cancer Cell. 35:347–367.

Cukierman, E., R. Pankov, D.R. Stevens, and K.M. Yamada. 2001. Taking cellmatrix adhesions to the third dimension. Science. 294:1708–1712.

Denk-Lobnig, M., and A.C. Martin. 2019. Modular regulation of Rho family GTPases in development. Small GTPases. 10:122–129.

Dozynkiewicz, M.A., N.B. Jamieson, I. Macpherson, J. Grindlay, P.V. van den Berghe, A. von Thun, J.P. Morton, C. Gourley, P. Timpson, C. Nixon, C.J. McKay, R. Carter, D. Strachan, K. Anderson, O.J. Sansom, P.T. Caswell, and J.C. Norman. 2012. Rab25 and CLIC3 collaborate to promote integrin recycling from late endosomes/lysosomes and drive cancer progression. Dev Cell. 22:131–145.

Egami, Y., T. Taguchi, M. Maekawa, H. Arai, and N. Araki. 2014. Small GTPases and phosphoinositides in the regulatory mechanisms of macropinosome formation and maturation. Front Physiol. 5:374.

Ferreira, A.P.A., and E. Boucrot. 2018. Mechanisms of Carrier Formation during clathrin-Independent Endocytosis. Trends Cell Biol. 28:188–200.

Fort, L., J.M. Batista, P.A. Thomason, H.J. Spence, J.A. Whitelaw, L. Tweedy, J. Greaves, K.J. Martin, K.I. Anderson, P. Brown, S. Lilla, M.P. Neilson, P. Tafelmeyer, S. Zanivan, S. Ismail, D.M. Bryant, N.C.O. Tomkinson, L.H. Chamberlain, G.S. Mastick, R.H. Insall, and L.M. Machesky. 2018. Fam49/CYRI interacts with Rac1 and locally suppresses protrusions. Nat Cell Biol. 20:1159–1171.

Fujii, M., K. Kawai, Y. Egami, and N. Araki. 2013. Dissecting the roles of Rac1 activation and deactivation in macropinocytosis using microscopic photomanipulation. Sci Rep. 3:2385.

Gu, Z., E.H. Noss, V.W. Hsu, and M.B. Brenner. 2011. Integrins traffic rapidly via circular dorsal ruffles and macropinocytosis during stimulated cell migration. J Cell Biol. 193:61–70.

Hayer, A., M. Stoeber, C. Bissig, and A. Helenius. 2010. Biogenesis of caveolae: stepwise assembly of large caveolin and cavin complexes. Traffic. 11:361–382.

Humphreys, D., V. Singh, and V. Koronakis. 2016. Inhibition of WAVE Regulatory Complex Activation by a Bacterial Virulence Effector Counteracts Pathogen Phagocytosis. Cell Rep. 17:697–707.

Journet, A., G. Klein, S. Brugiere, Y. Vandenbrouck, A. Chapel, S. Kieffer, C. Bruley, C. Masselon, and L. Aubry. 2012. Investigating the macropinocytic proteome of Dictyostelium amoebae by high-resolution mass spectrometry. Proteomics. 12:241–245.

Kaplan, E., R. Stone, P.J. Hume, N.P. Greene, and V. Koronakis. 2020. Structure of CYRI-B (FAM49B), a key regulator of cellular actin assembly. Acta Crystallogr D Struct Biol. 76:1015–1024.

Kavran, J.M., D.E. Klein, A. Lee, M. Falasca, S.J. Isakoff, E.Y. Skolnik, and M.A. Lemmon. 1998. Specificity and promiscuity in phosphoinositide binding by pleckstrin homology domains. J Biol Chem. 273:30497–30508.

Lai, C.L., A. Srivastava, C. Pilling, A.R. Chase, J.J. Falke, and G.A. Voth. 2013. Molecular mechanism of membrane binding of the GRP1 PH domain. J Mol Biol. 425:3073–3090.

Leslie, E.J., J.C. Carlson, J.R. Shaffer, E. Feingold, G. Wehby, C.A. Laurie, D. Jain, C.C. Laurie, K.F. Doheny, T. McHenry, J. Resick, C. Sanchez, J. Jacobs, B. Emanuele, A.R. Vieira, K. Neiswanger, A.C. Lidral, L.C. Valencia-Ramirez, A.M. Lopez-Palacio, D.R. Valencia, M. Arcos-Burgos, A.E. Czeizel, L.L. Field, C.D. Padilla, E.M. Cutiongco-de la Paz, F. Deleyiannis, K. Christensen, R.G. Munger, R.T. Lie, A. Wilcox, P.A. Romitti, E.E. Castilla, J.C. Mereb, F.A. Poletta, I.M. Orioli, F.M. Carvalho, J.T. Hecht, S.H. Blanton, C.J. Buxo, A. Butali, P.A. Mossey, W.L. Adeyemo, O. James, R.O. Braimah, B.S. Aregbesola, M.A. Eshete, F. Abate, M. Koruyucu, F. Seymen, L. Ma, J.E. de Salamanca, S.M. Weinberg, L. Moreno, J.C. Murray, and M.L. Marazita. 2016. A multi-ethnic genome-wide association study identifies novel loci for non-syndromic cleft lip with or without cleft palate on 2p24.2, 17q23 and 19q13. Hum Mol Genet. 25:2862–2872.

Makyio, H., M. Ohgi, T. Takei, S. Takahashi, H. Takatsu, Y. Katoh, A. Hanai, T. Ueda, Y. Kanaho, Y. Xie, H.W. Shin, H. Kamikubo, M. Kataoka, M. Kawasaki, R. Kato, S. Wakatsuki, and K. Nakayama. 2012. Structural basis for Arf6-MKLP1 complex formation on the Flemming body responsible for cytokinesis. EMBO J. 31:2590–2603.

Mierke, C.T., B. Frey, M. Fellner, M. Herrmann, and B. Fabry. 2011. Integrin alpha5beta1 facilitates cancer cell invasion through enhanced contractile forces. J Cell Sci. 124:369–383.

Millius, A., S.N. Dandekar, A.R. Houk, and O.D. Weiner. 2009. Neutrophils establish rapid and robust WAVE complex polarity in an actin-dependent fashion. Curr Biol. 19:253–259.

Mooren, O.L., B.J. Galletta, and J.A. Cooper. 2012. Roles for actin assembly in endocytosis. Annu Rev Biochem. 81:661–686.

Moreno-Layseca, P., N.Z. Jäntti, R. Godbole, C. Sommer, G. Jacquemet, H. Al-Akhrass, P. Kronqvist, R.E. Kallionpää, L. Oliveira-Ferrer, P. Cervero, S. Linder, M. Aepfelbacher, J. Rae, R.G. Parton, A. Disanza, G. Scita, S. Mayor, M. Selbach, S. Veltel, and J. Ivaska. 2020. Cargo-specific recruitment in clathrin and dynamin-independent endocytosis. BioRXIV.

Nam, J.M., Y. Onodera, M.J. Bissell, and C.C. Park. 2010. Breast cancer cells in three-dimensional culture display an enhanced radioresponse after coordinate targeting of integrin alpha5beta1 and fibronectin. Cancer Res. 70:5238–5248.

Paul, N.R., J.L. Allen, A. Chapman, M. Morlan-Mairal, E. Zindy, G. Jacquemet, L. Fernandez del Ama, N. Ferizovic, D.M. Green, J.D. Howe, E. Ehler, A. Hurlstone, and P.T. Caswell. 2015. alpha5beta1 integrin recycling promotes Arp2/3-independent cancer cell invasion via the formin FHOD3. J Cell Biol. 210:1013–1031.

Pietila, M., P. Sahgal, E. Peuhu, N.Z. Jantti, I. Paatero, E. Narva, H. Al-Akhrass, J. Lilja, M. Georgiadou, O.M. Andersen, A. Padzik, H. Sihto, H. Joensuu, M. Blomqvist, I. Saarinen, P.J. Bostrom, P. Taimen, and J. Ivaska. 2019. SORLA regulates endosomal trafficking and oncogenic fitness of HER2. Nat Commun. 10:2340.

Rainero, E., J.D. Howe, P.T. Caswell, N.B. Jamieson, K. Anderson, D.R. Critchley, L. Machesky, and J.C. Norman. 2015. Ligand-Occupied Integrin Internalization Links Nutrient Signaling to Invasive Migration. Cell Rep. 10:398–413.

Rizzo, M.A., M.W. Davidson, and D.W. Piston. 2009. Fluorescent protein tracking and detection: fluorescent protein structure and color variants. Cold Spring Harb Protoc. 2009:pdb.top 63.

Roy, A., A. Kucukural, and Y. Zhang. 2010. I-TASSER: a unified platform for automated protein structure and function prediction. Nat Protoc. 5:725–738.

Schlam, D., R.D. Bagshaw, S.A. Freeman, R.F. Collins, T. Pawson, G.D. Fairn, and S. Grinstein. 2015. Phosphoinositide 3-kinase enables phagocytosis of large particles by terminating actin assembly through Rac/Cdc42 GTPase-activating proteins. Nat Commun. 6:8623.

Schliwa, M. 1982. Action of cytochalasin D on cytoskeletal networks. J Cell Biol. 92:79–91.

Shang, W., Y. Jiang, M. Boettcher, K. Ding, M. Mollenauer, Z. Liu, X. Wen, C. Liu, P. Hao, S. Zhao, M.T. McManus, L. Wei, A. Weiss, and H. Wang. 2018. Genome-wide CRISPR screen identifies FAM49B as a key regulator of actin dynamics and T cell activation. Proc Natl Acad Sci U S A. 115:E4051–E4060.

Shi, F., and J. Sottile. 2008. Caveolin-1-dependent beta1 integrin endocytosis is a critical regulator of fibronectin turnover. J Cell Sci. 121:2360–2371.

Swanson, J.A., and C. Watts. 1995. Macropinocytosis. Trends Cell Biol. 5:424–428.

Timpson, P., E.J. McGhee, Z. Erami, M. Nobis, J.A. Quinn, M. Edward, and K.I. Anderson. 2011. Organotypic collagen I assay: a malleable platform to assess cell behaviour in a 3-dimensional context. J Vis Exp:e3089.

Varnai, P., and T. Balla. 1998. Visualization of phosphoinositides that bind pleckstrin homology domains: calcium-and agonist-induced dynamic changes and relationship to myo-[3H]inositol-labeled phosphoinositide pools. J Cell Biol. 143:501–510.

Varnai, P., T. Bondeva, P. Tamas, B. Toth, L. Buday, L. Hunyady, and T. Balla. 2005. Selective cellular effects of overexpressed pleckstrin-homology domains that recognize PtdIns(3,4,5)P3 suggest their interaction with protein binding partners. J Cell Sci. 118:4879–4888.

Veltman, D.M., T.D. Williams, G. Bloomfield, B.C. Chen, E. Betzig, R.H. Insall, and R.R. Kay. 2016. A plasma membrane template for macropinocytic cups. Elife. 5.

Yang, J., R. Yan, A. Roy, D. Xu, J. Poisson, and Y. Zhang. 2015. The I-TASSER Suite: protein structure and function prediction. Nat Methods. 12:7–8.

Yarmola, E.G., T. Somasundaram, T.A. Boring, I. Spector, and M.R. Bubb. 2000. Actin-latrunculin A structure and function. Differential modulation of actin-binding protein function by latrunculin A. J Biol Chem. 275:28120–28127.

Yelland, T., A.H. Le, S. Nikolaou, R. Insall, L. Machesky, and S. Ismail. 2020. Structural Basis of CYRI-B Direct Competition with Scar/WAVE Complex for Rac1. Structure.

Yoshida, S., A.D. Hoppe, N. Araki, and J.A. Swanson. 2009. Sequential signaling in plasma-membrane domains during macropinosome formation in macrophages. J Cell Sci. 122:3250–3261.

Yu, X., T. Zech, L. McDonald, E.G. Gonzalez, A. Li, I. Macpherson, J.P. Schwarz, H. Spence, K. Futo, P. Timpson, C. Nixon, Y. Ma, I.M. Anton, B. Visegrady, R.H. Insall, K. Oien, K. Blyth, J.C. Norman, and L.M. Machesky. 2012. N-WASP coordinates the delivery and F-actin-mediated capture of MT1-MMP at invasive pseudopods. J Cell Biol. 199:527–544.

Yuki, K.E., H. Marei, E. Fiskin, M.M. Eva, A.A. Gopal, J.A. Schwartzentruber, J. Majewski, M. Cellier, J.N. Mandl, S.M. Vidal, D. Malo, and I. Dikic. 2019. CYRI/FAM49B negatively regulates RAC1-driven cytoskeletal remodelling and protects against bacterial infection. Nat Microbiol. 4:1516–1531.

Zech, T., S.D. Calaminus, P. Caswell, H.J. Spence, M. Carnell, R.H. Insall, J. Norman, and L.M. Machesky. 2011. The Arp2/3 activator WASH regulates alpha5beta1-integrin-mediated invasive migration. J Cell Sci. 124:3753–3759.

Zhang, Y. 2008. I-TASSER server for protein 3D structure prediction. BMC Bioinformatics. 9:40.

